# Cholesterol deficiency directs autophagy-dependent secretion of extracellular vesicles

**DOI:** 10.1101/2025.01.11.632510

**Authors:** Jazmine D. W. Yaeger, Sonali Sengupta, Austin L. Walz, Mayu Morita, Terry K. Morgan, Paola D. Vermeer, Kevin R. Francis

**Author notes:** Corresponding author (K. R. F.).

## Abstract

Extracellular vesicle (EV) secretion is an important, though not fully understood, intercellular communication process. Lipid metabolism has been shown to regulate EV activity, though the impact of specific lipid classes is unclear. Through analysis of small EVs (sEVs), we observe aberrant increases in sEV release within genetic models of cholesterol biosynthesis disorders, where cellular cholesterol is diminished. Inhibition of cholesterol synthesis at multiple synthetic steps mimics genetic models in terms of cholesterol reduction and sEVs secreted. Further analyses of sEVs from cholesterol-depleted cells revealed structural deficits and altered surface marker expression, though these sEVs were also more easily internalized by recipient cells. Transmission electron microscopy of cells with impaired cholesterol biosynthesis demonstrated multivesicular and multilamellar structures potentially associated with autophagic defects. We further found autophagic vesicles being redirected toward late endosomes at the expense of autophagolysosomes. Through CRISPR-mediated inhibition of autophagosome formation, we mechanistically determined that release of sEVs after cholesterol depletion is autophagy dependent. We conclude that cholesterol imbalance initiates autophagosome-dependent secretion of sEVs, which may have pathological relevance in diseases of cholesterol disequilibrium.

## INTRODUCTION

Cholesterol is a vital component of cell membranes, providing structural integrity to organelles, anchoring proteins and carbohydrates to subcellular spaces, and serving as a precursor in steroidogenesis. In this way, cholesterol metabolism functions to both directly and indirectly facilitate critical cellular processes. These lipid-influenced regulatory pathways include cellular trafficking in the forms of endosomal sorting, extracellular transport, and the underexplored area of extracellular vesicle (EV) formation and release (Meldolesi, 2018; Record et al., 2018). While EV function and cargo receive attention due to their close association with pathological states such as cancer (Hoshino et al., 2020; Vinduska et al., 2021), the mechanisms regulating EV biosynthesis and function require investigation.

A broad range of EVs have been categorized based on size and origin (Buzas, 2022; Meldolesi, 2018; Van der Pol et al., 2012), including apoptotic bodies (>1000 nm diameter) and microvesicles (300-1000 nm diameter) which are derived from membrane blebbing. However, small EVs (sEVs; 50-200 nm diameter), sometimes called exosomes, emerge from a complex and tightly regulated intracellular process. Beginning with invaginations of early endosomes called intraluminal vesicles (ILVs) then packaged into multivesicular bodies (MVBs), this process is inseparable from endosomal sorting pathways. The fate of MVBs is not linear, as they may merge with lysosomes (endolysosome formation) for degradation, localize to the plasma membrane for extracellular release, or fuse with autophagy-related endosomes to form autolysosomes or amphisomes (Ganesan and Cai, 2021; van Niel et al., 2018).

Compared to the cell of origin, sEVs are reportedly enriched in cholesterol (Skotland et al., 2020). Cholesterol is selectively sorted into MVBs via the tetraspanin and sEV surface marker CD63 (Palmulli et al., 2024). Blockade of early cholesterol biosynthesis through statin-mediated inhibition of HMG-CoA reductase (HMGCR) was shown to reduce expression of common sEV markers (Kulshreshtha et al., 2019; Sekhavati et al., 2023). Additionally, studies investigating the cholesterol storage disorder Niemann-Pick disease, Type C1 suggested aberrantly accumulating cholesterol is released in sEVs (Ilnytska et al., 2021; Strauss et al., 2010). While these findings suggest a relationship between cholesterol and sEV biology, the enigmatic association between sEV dynamics and cholesterol metabolism remains unresolved.

Cholesterol biosynthesis begins when acetyl-CoA enters the mevalonate pathway. After a series of enzymatic steps, including the conversion of HMG-CoA to mevalonate via the catalytic activity of HMGCR, squalene is formed and sterol biosynthesis may proceed through the Kandutsch-Russell or Bloch pathways. Disruption of cholesterol metabolism can result in both reduced cholesterol production and accumulation of other sterols. Importantly, cholesterol disequilibrium is commonly associated with numerous diseases, including cancer and neurodegenerative disorders, like Alzheimer’s disease (Dai et al., 2021; Mayengbam et al., 2021; Xue-shan et al., 2016), which may be consequentially tied to changes in sEV production observed in the same pathological conditions (Clayton et al., 2021; Pegtel et al., 2014; Soria et al., 2017). Disorders of cholesterol metabolism also include rare genetic disorders due to mutations within cholesterol synthetic genes. The most common of these disorders, Smith-Lemli-Opitz syndrome (SLOS), occurs when *DHCR7*, the gene coding for 7-dehydrocholesterol reductase, is mutated leading to decreased cellular cholesterol and accumulation of the sterol 7-dehydrocholesterol (7-DHC) (Fitzky et al., 1998; Wassif et al., 1998). Other less common genetic disruptions affecting late-stage cholesterol biosynthesis result in additional rare disorders including lathosterolosis (LATH; *SC5D* mutation) and desmosterolosis (*DHCR24* mutation) (Krakowiak et al., 2003; Schaaf et al., 2011). We have previously shown that disruption of cholesterol homeostasis and an increase in sterol precursors impairs clathrin-mediated endocytosis and membrane bending (Anderson et al., 2021), suggesting sterol-impacted membrane may disrupt sEV production, packaging, or utility.

In this study, we determine cholesterol biosynthesis is an important regulator of sEV formation, phenotype, and function. Cellular models of cholesterol biosynthesis disorders, SLOS and LATH, secrete more sEVs relative to controls. We investigated this phenotype further using pharmacological inhibition of cholesterol metabolism at varying steps in the pathway, observing enhanced sEV release across cell types upon cholesterol disruption, an effect strongly correlated to cellular cholesterol content. Additionally, we find phenotypic and structural alterations in cholesterol-disrupted sEVs that impact internalization, as well as stalled endosomal sorting pathways. We lastly determine autophagy is a critical component for sEV release in cholesterol-depleted conditions. These discoveries mechanistically link cholesterol metabolism, lipid transport, and sEV biogenesis, detailing disorganization of cellular trafficking mechanisms with high relevance to pathologies exhibiting loss of cholesterol homeostasis.

## RESULTS

### Cellular cholesterol regulates small extracellular vesicle release

To determine if sEV production and release are impacted in diseases of cholesterol metabolism, we first assessed sEV secretion in human fibroblasts derived from SLOS and LATH subjects (**Figure 1A**). Following EV depletion from fetal bovine serum (FBS) and cholesterol-depleted lipoprotein deficient serum (LPDS) (**Figure S1A**), control and patient-derived fibroblasts were analyzed for biochemical deficits and sEV release (**Figure S1B**). Using gas chromatography with mass spectrometry (GC-MS), we observed SLOS and LATH fibroblasts exhibit reduced cholesterol levels compared to controls, with accumulation of sterol precursor molecules in SLOS and LATH samples (7-DHC and lathosterol, respectively) (**Figure 1B, Figure S2A**). We then purified sEVs by differential centrifugation and analyzed them using nanoparticle tracking analysis (NTA) (**Figure S1B**). sEV numbers were increased in SLOS and LATH fibroblasts relative to controls (**Figure 1C, Figure S2B, C**), suggesting cholesterol homeostasis is an important regulator of sEV biology.

**Figure 1.**
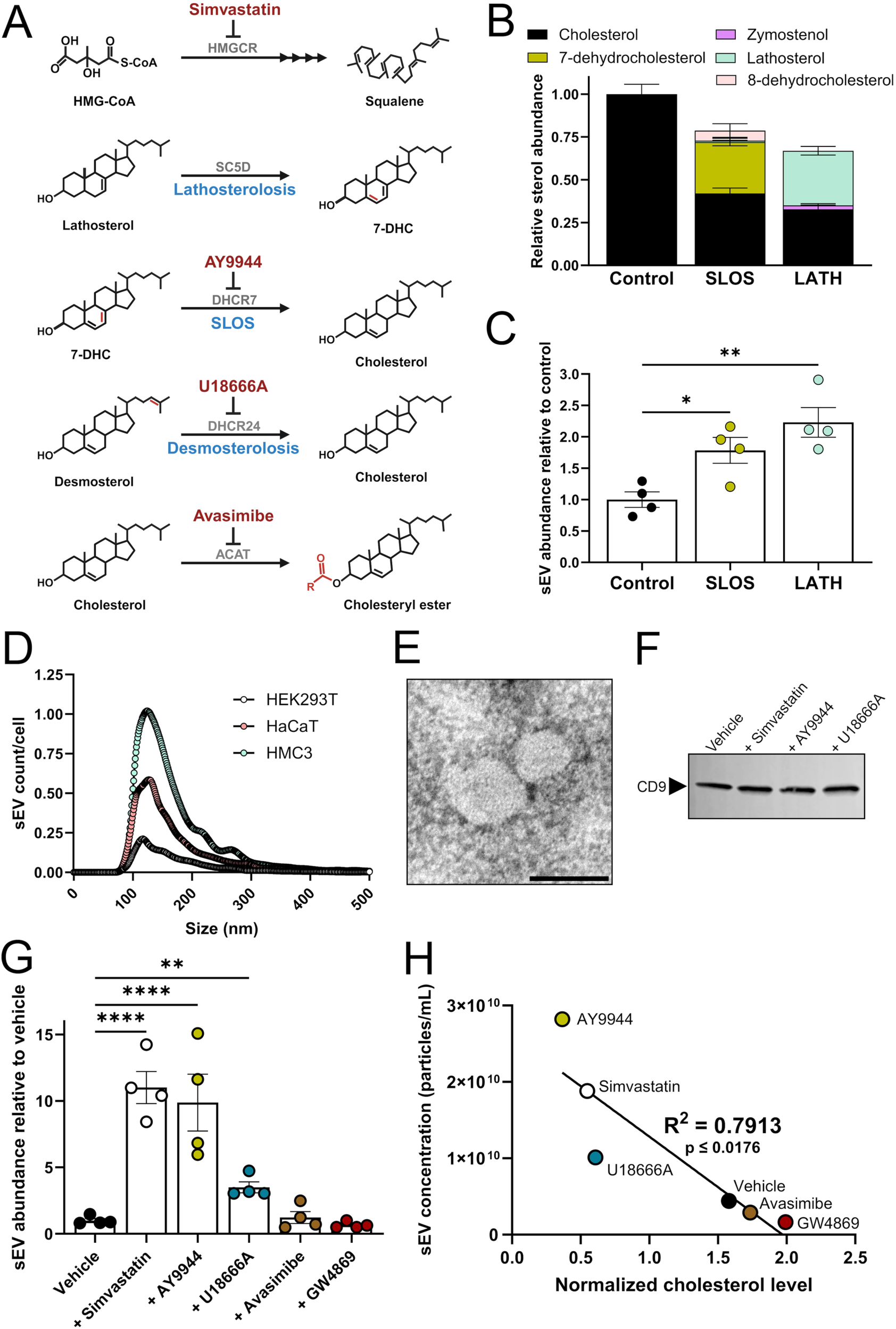
Cholesterol depletion promotes small extracellular vesicle release. (**A**) Cholesterol metabolic reactions targeted with small molecule inhibitors (red text). Rare diseases of cholesterol biosynthesis (blue text) arise from genetic mutations to indicated enzymes (gray). (**B**) GC-MS analysis of human fibroblasts shows Smith-Lemli-Opitz syndrome (SLOS) and lathosterolosis (LATH) are cholesterol deficient and accumulate sterol intermediates (mean ± SEM; n = 3 biological replicates from 3 independent experiments). (**C**) Small extracellular vesicle (sEV) release is elevated in SLOS and LATH fibroblasts relative to controls (mean ± SEM; n = 4 biological replicates from 4 independent experiments). One-way ANOVA (F2,9 = 10.22, p ≤ 0.0048), Dunnett’s multiple comparisons; *p ≤ 0.05; **p ≤ 0.005. (**D**) Nanoparticle tracking analysis (NTA) of sEV size from HEK293T, HaCaT, and HMC3 cells (mean; n = 4 biological replicates from 4 independent experiments). (**E**) Representative transmission electron microscopy (TEM) image of sEVs isolated from HEK293T cells. Scale bar, 100 nm. (**F**) CD9 immunoblot in HEK293T-isolated sEVs following treatment with respective cholesterol biosynthesis inhibition. (**G**)Inhibition of cholesterol biosynthesis in HEK293T cells results in increased sEV release (mean ± SEM; n = 4 biological replicates from 4 independent experiments). One-way ANOVA (F5,18 = 20.51, p < 0.0001), Dunnett’s multiple comparisons test; **p ≤ 0.005, ****p < 0.0001). (**H**) Regression analysis of sEVs from HEK293T cells reveals a strong negative correlation between sEV secretion and cellular cholesterol levels. Linear regression (F1,4 = 15.17, R2 = 0.7913, p ≤ 0.0176).

To more thoroughly investigate the relationship between sEV release and cholesterol metabolism, we treated three human cell models, embryonic kidney cells (HEK293T), keratinocytes (HaCaT), and microglia (HMC3), with small molecule inhibitors of cholesterol biosynthesis. Simvastatin, AY9944, and U18666A reduced cellular cholesterol in all three cell types (**Figure S3**). As simvastatin impairs cholesterol biosynthesis early in the pathway, prior to sterol production (**Figure 1A**), lower cholesterol levels are observed in the absence of precursor sterol accumulation. Alternatively, AY9944 inhibition of DHCR7 (**Figure 1A**) both reduces cholesterol levels and causes a buildup of other sterols, predominantly 7-DHC, zymostenol, and 7,24-cholestadienol in all cell types (**Figure S3**). Antagonism of DHCR24 with the small molecule U18666A (**Figure 1A**) reduces cholesterol levels most strongly in HaCaT cells and further leads to accumulation of desmosterol (**Figure S3**). While neither the acyl-coA:cholesterol acyltransferase (ACAT) inhibitor avasimibe nor the neutral sphingomyelinase (nSMase) antagonist GW4869 altered the cellular levels of cholesterol, we observed accumulation of desmosterol with GW4869 treatment in all cell types (**Figure S3**). These results demonstrate small molecule inhibitors efficiently alter sterol profiles in these distinct cell types, modeling disease biochemical phenotypes and allowing us to discern cell type dependent impacts on sEV biology.

To examine the effect of cellular sterol content on sEV production, we quantified secreted sEVs within the media after treatment with small molecule inhibitors. At baseline, the particles secreted by HEK293T, HaCaT, and HMC3 cells were within the 75 nm - 300 nm size range, with the highest concentration being between 100 nm – 200 nm (**Figure 1D**). When normalized by cell count, HMC3 cells produced the greatest number of particles, followed by HaCaT and HEK293T cells, respectively (**Figure 1D**). Beyond NTA of particle number and size, sEV isolations were also validated using transmission electron microscopy (TEM) (**Figure 1E**) and immunoblotting for the common sEV tetraspanin CD9 (**Figure 1F**).

Similar to SLOS and LATH fibroblasts, pharmacological blockade of cholesterol biosynthesis for 48 h increased sEV release in all three cell lines (**Figure 1G, Figure S4A-C**). Avasimibe or GW4869 treatments did not significantly impact the number of secreted vesicles (**Figure 1G, Figure S4A, B, C**). To further define the relationship between cholesterol homeostasis and sEV release, we performed regression analyses for HEK293T, HaCaT, and HMC3 cells treated with small molecule inhibitors. In each cell type, we observed a negative correlation between cellular cholesterol and sEVs released (**Figure 1H, Figure S4D, E**). To determine if increased sEV secretion in response to cholesterol depletion was time dependent, we assessed sEV release after 24 h of treatment. While the overall number of sEVs were reduced, we observed a similar increase in sEV secretion following cholesterol disruption (**Figure S5A**). Importantly, although these treatments impaired cell proliferation (**Figure S5B, C, Figure S6A, B**), increased sEV release due to cholesterol synthesis inhibition was not driven by apoptosis and production of apoptotic bodies (**Figure S6C**). Collectively, these findings demonstrate intracellular cholesterol content is directly correlative to sEV production and secretion.

### Cholesterol depletion phenotypically modifies EV populations

To better understand the observed changes in sEV output after blockade of cholesterol biosynthesis, we analyzed the phenotype of secreted sEVs. First, we assessed sEV surface marker expression using a MACSplex fluorescent bead-based assay, utilizing flow cytometry to quantify distinct populations of sEVs based on surface protein expression (**Figure 2A; Table S1**). Cholesterol synthesis inhibitors exhibited drug-specific impacts on sEV profiles. While all treatment groups expressed high levels of CD9, CD63, and CD81 tetraspanin proteins, sEVs from AY9944-treated cells exhibited reduced levels of CD63 and CD81 relative to controls (**Figure 2B, D**). Meanwhile, simvastatin and U18666A treatments resulted in elevated expression of CD56, CD29, and ROR1 on secreted sEVs, while AY9944 treatment produced sEVs rich in CD3, CD56, CD146, and ROR1 (**Figure 2B-D**). U18666A also produced sEVs expressing increased CD146, CD24, CD326, and CD29 relative to vehicle (**Figure 2B, C**). Expression patterns for CD29 and ROR1 in sEVs were further validated by immunoblot, where we observed increased expression of both proteins in all cholesterol biosynthesis inhibitor treatment groups relative to vehicle (**Figure 2E**). While avasimibe-treated sEVs exhibited decreased expression of CD3, MCSP, and CD29 relative to vehicle (**Figure 2B, C**), sEVs secreted from GW4869-treated cells were similar to controls (**Figure 2B, C, Table S1**). These results suggest cholesterol depletion alters the signaling capacity of sEVs in a biochemical-specific manner.

**Figure 2.**
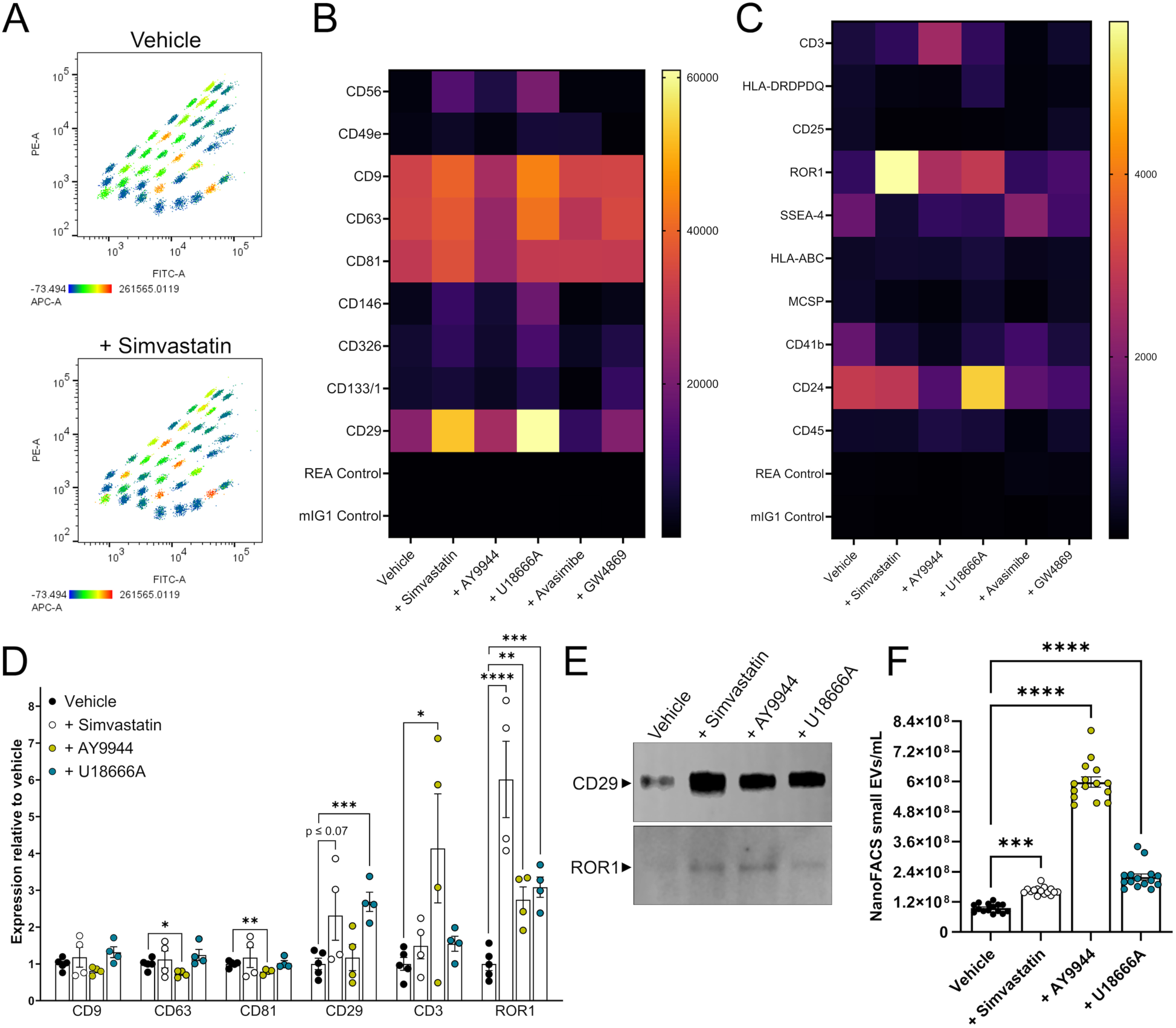
Cholesterol depletion alters sEV surface epitope expression and sEV pools. (**A**) Representative plots illustrating gating strategy for sEV epitope expression. Clustered populations represent expression of individual epitopes on sEVs. (**B**) Heat map representation of intensity values for highly expressed sEV epitopes following drug treatment (median; n = 4 biological replicates from 4 independent experiments). (**C**) Heat map representation of intensity values for moderately expressed sEV epitopes following drug treatment (median; n = 4 biological replicates from 4 independent experiments). (**D**) Quantification of cholesterol depletion impacts on selected sEV epitopes (mean ± SEM; n = 4 biological replicates from 4 independent experiments). One-way ANOVA (CD63: F3,13 = 4.714, p ≤ 0.0194; CD81: F3,13 = 20.52, p < 0.0001; CD29: F3,13 = 6.801, p ≤ 0.0054; CD3: F3,13 = 10.12, p ≤ 0.001; ROR1: F3,13 = 24.01, p < 0.0001). Unpaired t-test (CD63: t7 = 2.937, *p ≤ 0.05; CD81: t7 = 4.07, **p ≤ 0.005; CD29: Vehicle vs +Simvastatin, t7 = 2.135, p ≤ 0.0702, Vehicle vs +U18666A, t7 = 5.832, ***p ≤ 0.001; CD3: t7 = 2.384, *p ≤ 0.05; ROR1: t7 = 4.719, **p ≤ 0.005). (**E**) Expression of CD29 and ROR1 in isolated sEVs subsequent to cholesterol depletion. (**F**) NanoFACS analyses of small (<200 nm) EVs measured from media of HEK293T cells after cholesterol biosynthesis inhibition (mean ± SEM; n = 5 technical replicates from 3 biological replicates from 3 independent experiments). One-way ANOVA (F3,56 = 331.9, p < 0.0001). Dunnett’s multiple comparisons test (***p ≤ 0.001; ****p < 0.0001).

To further validate sEV phenotypes resulting from our cholesterol-depleted conditions, we used nano-flow cytometry (NanoFACS) to measure EVs directly from media without the need for differential centrifugation isolation (**Figure 2F, Figure S7A**). NanoFACS quantitatively measures EV size to validate that the increased sEV numbers observed were not due to breakdown of larger vesicles during standard isolation techniques (Morgan, 2018). As with our standard centrifugation isolations, NanoFACS demonstrated that simvastatin, AY9944, and U18666A increased sEV release relative to vehicle treatment (**Figure 2F**), while vehicle-treated cells produced increased numbers of large EVs (200 nm – 900 nm) compared to cholesterol-depleted conditions (**Figure S7B**). Importantly, the number of large events measured was orders of magnitude less than that of sEVs in all treatment groups (**Figures 2F, Figure S7B**). These experiments demonstrate cholesterol depletion directs cells to secrete more smaller vesicles, likely exosomes or exosome-like structures.

To determine if cholesterol deficient conditions impacted the physical structure of sEVs, we analyzed sEV morphology by TEM following cholesterol synthesis inhibition (**Figure 3A-C**). Size analyses showed vehicle-treated sEVs generally fell between 60-120 nm, while simvastatin or AY9944 treatment produced sEVs more evenly distributed from 60 nm to >200 nm (**Figure 3B**). The relative frequency of spherical sEVs, determined by aspect ratio (major/minor diameter), was reduced in simvastatin or AY9944 treatments (**Figure 3C**). These data demonstrate cholesterol homeostasis is critical to maintenance of sEV shape and structure. Since an important biological function of cholesterol is to provide structural support to membrane-bound compartments, we predicted sEV cholesterol levels would be proportionally decreased following cholesterol depletion. GC-MS analysis of sterol abundance after simvastatin, AY9944, or U18666A treatments revealed an exaggerated biochemical deficiency compared to vehicle (**Figure 3D**). sEV cholesterol deficiency was significantly and positively correlated to the cholesterol levels in secreting cells (**Figure 3E**). These results demonstrate cholesterol-depleted conditions produce structurally aberrant sEVs that likely exhibit modified signaling capacity.

**Figure 3.**
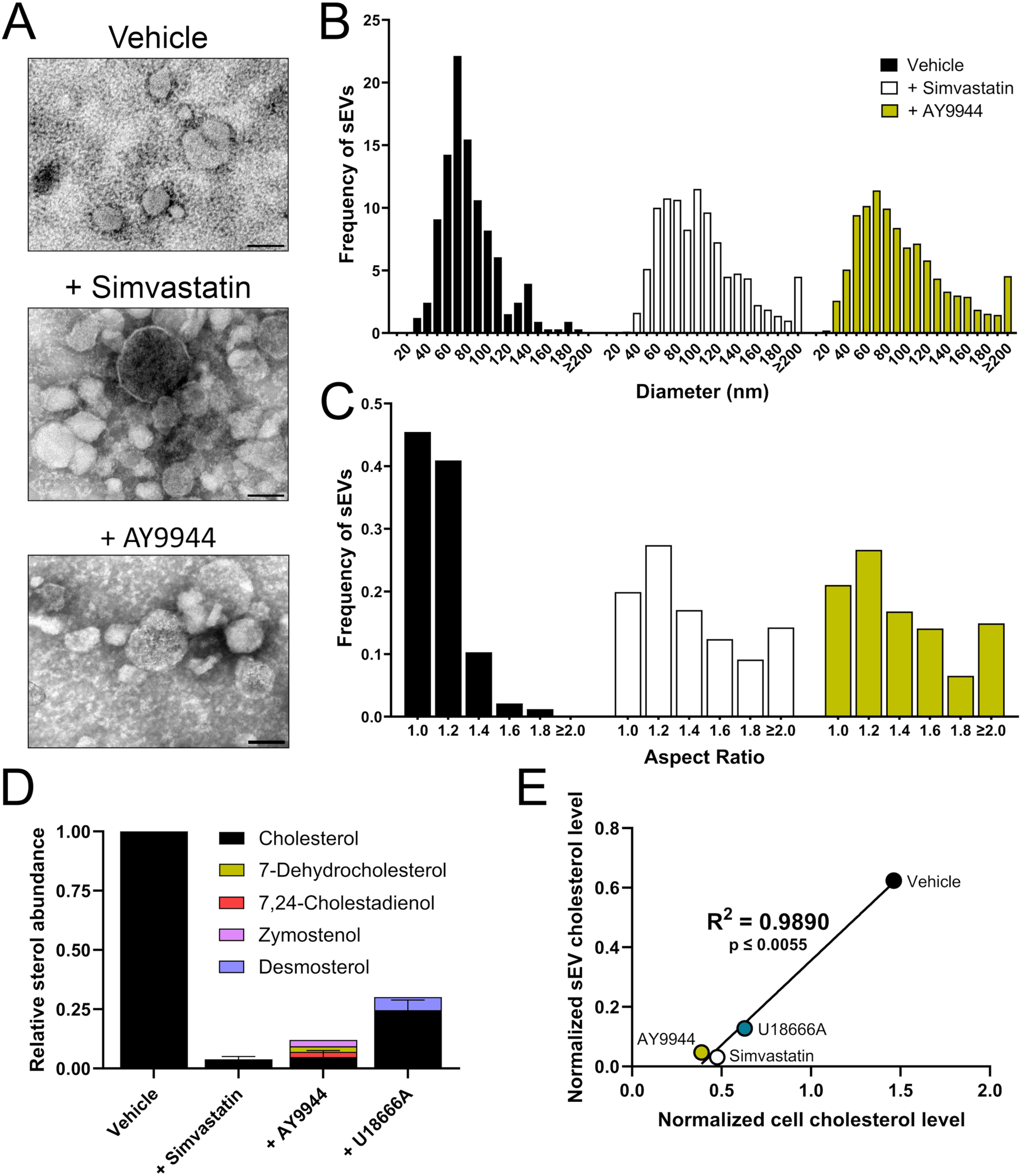
sEVs from cholesterol-depleted cells are structurally aberrant. (**A**) Representative TEM images of sEVs derived from vehicle and cholesterol-targeted HEK293T cells. Scale bar, 100 nm. (**B**) sEV size distribution after cholesterol depletion drug treatment in HEK293T cells. (**C**) Spherical sEV structure analyzed by aspect ratio (major/minor width) demonstrates sEV asymmetry upon cholesterol depletion. (**D**) GC-MS analysis of sterol content in sEVs collected from cholesterol-depleted HEK293T cells (mean ± SEM; n = 2 biological replicates from 2 independent experiments). (**E**) Cellular cholesterol levels positively correlate with sEV cholesterol levels. Linear regression (F1,2 = 179.8, p ≤ 0.0055).

### Cholesterol-depleted cells generate sEVs that are more easily internalized

As cholesterol biosynthesis antagonism produced phenotypically and structurally aberrant sEVs, we asked if sEVs from cholesterol-inhibited cells could functionally internalize. We thus generated cells which stably overexpress fluorescently labeled sEV markers CD9 (HEK293T^Emerald-CD9^) or CD63 (HEK293T^eGFP-CD63^), followed by treatment with cholesterol biosynthesis inhibitors (**Figure 4A**). sEVs were isolated and quantified from HEK293T^Emerald-CD9^ and HEK293T^eGFP-CD63^ cells treated with vehicle, simvastatin, or AY9944 before incubating with untreated, recipient HEK293T (**Figure 4B**). Following labeling of the recipient cell plasma membrane (594-conjugated concanavalin A) and nucleus (Hoechst), sEV uptake was quantified (**Figure 4B-F**). 3D renderings from optically dissected z-stack images revealed fluorescently labeled sEVs were internalized and not adhered to the cell surface (**Figure 4C**). Both mEmerald-CD9-positive (**Figure 4D, E, Figure S8**) and eGFP-CD63-positive sEVs (**Figure 4F**) from simvastatin or AY9944-treated cells were more readily internalized compared to vehicle-treated in a dose-dependent manner. These findings demonstrate the signaling and structural changes induced by cholesterol-deficient conditions boosts sEV uptake by recipient cells.

**Figure 4.**
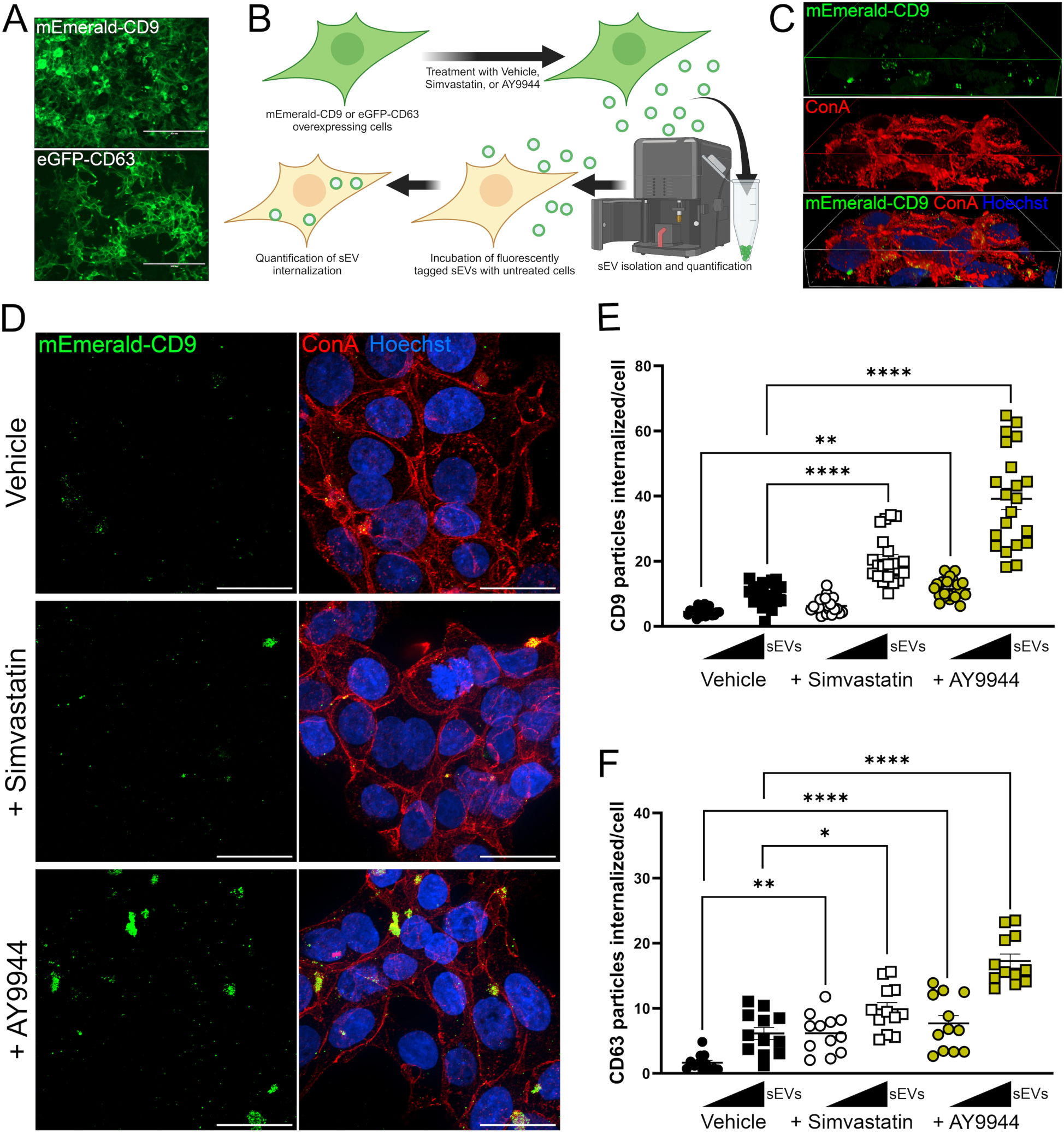
sEVs from cholesterol-depleted cells are more readily internalized. (**A**) Stably transfected HEK293T cells overexpressing mEmerald-CD9 or eGFP-CD63. Scale bar, 200 µm. (**B**) Experimental design for isolation of mEmerald-CD9 and eGFP-CD63 sEVs and quantitation of uptake by recipient cells. (**C**) Representative 3D reconstructions of target cell-uptake of mEmerald-CD9 sEVs. Plasma membrane visualized with concanavalin A (ConA). (**D**) Representative images of target cells incubated with sEVs from cells treated with vehicle, simvastatin, or AY9944. Scale bar, 25 µm. (**E**) Quantified mEmerald-CD9 sEV uptake in recipient cells (mean ± SEM; n = 21, 7 images analyzed from 3 independent experiments). Two-way ANOVA (Interaction effect: F2,120 = 26.31, p < 0.0001; Treatment effect: F2,120 = 70.3, p < 0.0001; Dose effect: F1,120 = 152.1, p < 0.0001). Sidak’s multiple comparisons test (**p ≤ 0.005, ****p < 0.0001). (**F**) Quantified eGFP-CD63 sEVs uptake in recipient cells (mean ± SEM; n = 12, 4 images analyzed from 3 independent experiments). Two-way ANOVA (Interaction effect: F2,66 = 5.633, p ≤ 0.0055; Treatment effect: F2,66 = 40.73, p < 0.0001; Dose effect: F1,66 = 58.16, p < 0.0001). Sidak’s multiple comparisons test (*p ≤ 0.05, **p ≤ 0.005, ****p < 0.0001).

### Blockade in endosomal sorting pathways leads to vesicle buildup after cholesterol starvation

As sEVs are derived from the same intracellular mechanisms that govern endosome formation and vesicular transport (Arya et al., 2024), we examined the impact of cholesterol synthesis inhibition on endosomal vesicle content. AY9944-treated cells exhibited increased expression of early endosomes, characterized by GTPase, Ras-related protein Rab5 (Rab5) expression (**Figure 5A, C**). Analysis of the late endosome marker, Rab7, revealed a significant increase in expression after AY9944 treatment (**Figure 5B, D**). Expression of the late endosome/sEV marker CD63 was also elevated in simvastatin and AY9944-treated cells (**Figure 5E, Figure S9A**). Assessment of the lysosomal marker lysosomal-associated protein 1 (LAMP1) showed perinuclear staining in all treatments (**Figure S9B, C**), a common impact of nutrient starvation (Korolchuk et al., 2011; Lee et al., 2020). To better visualize intracellular changes in endosomal pathways, we next utilized TEM (**Figure 5F**). Vehicle-treated cells revealed large mitochondria with few endosomal vesicles; however, simvastatin treatment produced small mitochondria and misshapen endosomal vesicles in the form of multilamellar bodies (**Figure 5F**). Similarly, AY9944 treatment produced small mitochondria and large, double membrane structures encompassing several multilamellar vesicles (**Figure 5F**). These findings suggest defects in endosomal sorting resulting from cholesterol depletion may be linked to sEV malformations.

**Figure 5.**
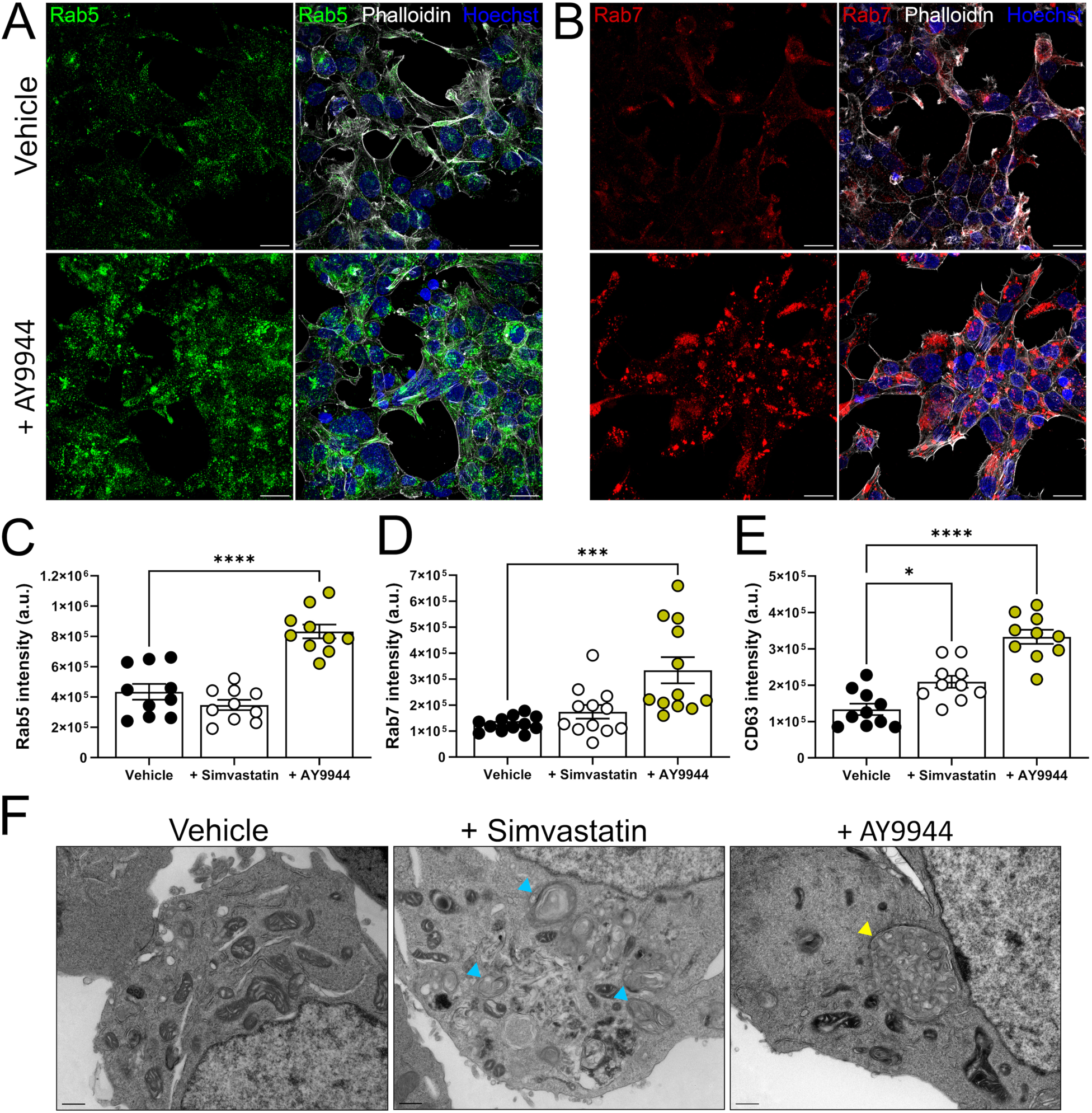
Cholesterol depletion causes defects in endosomal sorting and accumulation of multilamellar structures. (**A**) Representative images of Rab5 expression in HEK293T cells. Scale bar, 25 µm. (**B**) Representative images of Rab7 expression in HEK293T cells. Scale bar, 25 µm. (**C**) Cholesterol depletion causes increased expression of Rab5 (mean ± SEM; n = 10 images analyzed from 2 independent experiments). One-way ANOVA (F2,27 = 33.97, p < 0.0001), Dunnett’s multiple comparisons test; ****p < 0.0001. (**D**) Rab7 expression is increased with cholesterol depletion (n = 12 images analyzed from 3 independent experiments). One-way ANOVA (F2,33 = 10.7, p ≤ 0.0003), Dunnett’s multiple comparisons test; ***p ≤ 0.0003. (**E**) CD63 expression is increased with cholesterol depletion treatments (mean ± SEM; n = 10 images analyzed from 2 independent experiments). One-way ANOVA (F2,27 = 33.88, p < 0.0001), Dunnett’s multiple comparisons test; **p ≤ 0.0086, ****p < 0.0001. (**F**) TEM images of HEK293T cells treated with simvastatin or AY9944 reveal multilamellar structures (blue indicators) and multilamellar, multivesicular bodies (yellow indicators). Scale bar, 500 nm.

### Cholesterol biosynthesis inhibition causes autophagosome redirection to late endosomes

The appearance of multi-membraned structures upon cholesterol deficiency suggested that autophagosome pathways may be altered by cholesterol-targeted treatments. Macroautophagy, often referred to simply as autophagy, is stimulated during periods of cellular stress such as nutrient starvation (Mizushima and Komatsu, 2011). Importantly, the context surrounding autophagy induction may direct autophagic targets (Li et al., 2021), a mechanism of specificity not yet fully understood. In nutrient rich conditions, mature autophagosomes fuse with lysosomes to form an autolysosome. Under conditions of environmental stress, autophagosomes may be directed toward endosomal vesicles, creating amphisomes (**Figure 6A**). Analysis of the mature autophagosome marker LC3B showed simvastatin and AY9944 enhanced LC3B expression compared to vehicle treatment (**Figure 6B, C**). These data suggest cholesterol-deficient cells accumulate autophagic vesicles.

**Figure 6.**
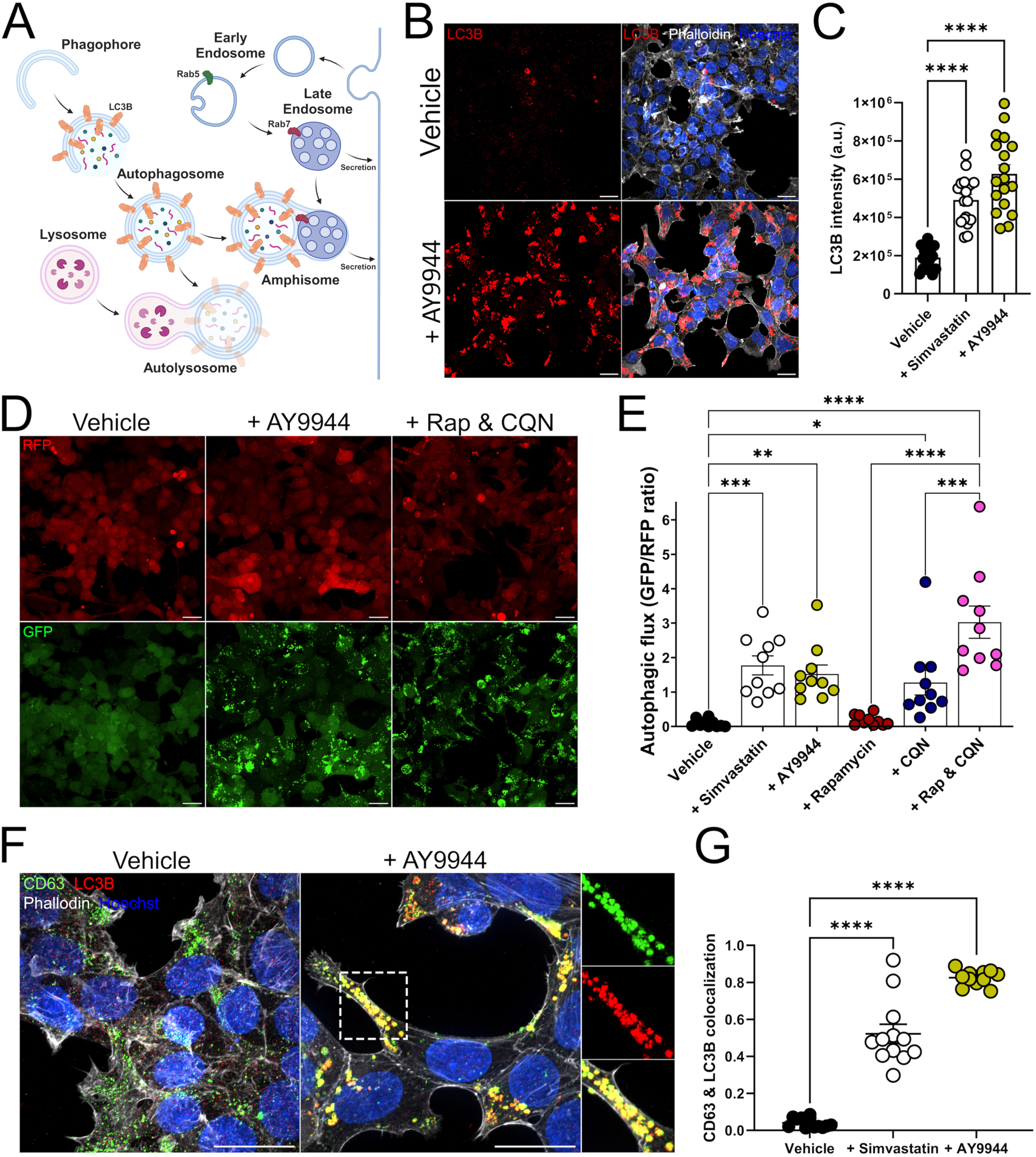
Cholesterol depletion induces dysregulation of autophagy-associated signaling mechanisms. (**A**) Illustration of normal autophagic signaling. Mature autophagosomes associate with lysosomes to form autolysosomes. If autolysosome formation is impaired, autophagosome/late endosome fusion forms amphisomes. (**B**) Representative images of LC3B expression in vehicle or AY9944 treated HEK293T cells. Scale bar, 25 µm. (**C**) Analysis of LC3B expression with cholesterol depletion (mean ± SEM; n = 18, 9 images analyzed from 3 independent experiments). One-way ANOVA (F2,51 = 47.57, p < 0.0001), Dunnett’s multiple comparisons test; ****p < 0.0001. (**D**) Representative images from HEK293TGFP-LC3-RFP-LC3ΔG cells treated with vehicle, AY9944, or rapamycin and chloroquine (Rap & CQN). Scale bar, 25 µm. (**E**) Analysis of autophagic flux upon cholesterol depletion or Rap & CQN (mean ± SEM; n = 10, 5 images analyzed from 2 independent experiments). One-way ANOVA (F5,54 = 14.51, p < 0.0001). Sidak’s multiple comparisons test (*p ≤ 0.05, **p ≤ 0.01, ***p ≤ 0.001, ****p < 0.0001). (**F**) Representative images of CD63 and LC3B expression in vehicle or AY9944-treated HEK293T cells. Scale bar, 25 µm. (**G**) Analyses reveal significant colocalization between CD63 and LC3B upon cholesterol depletion (mean ± SEM; n = 12 images analyzed from 3 independent experiments). One-way ANOVA (F2,33 = 164.8, p < 0.0001), Dunnett’s multiple comparisons test; ****p < 0.0001.

As accumulation of autophagosomes could be due to either increased autophagic signaling or limited autophagosome turnover, we generated a HEK293T line expressing a dual fluorescence, autophagic flux reporter (HEK293T^GFP-LC3-RFP-LC3ΔG^) to measure autophagosome turnover (Kaizuka et al., 2016). Dual treatment with the mammalian target of rapamycin (mTOR) inhibitor rapamycin to promote autophagosome formation and chloroquine (CQN) to prevent autophagosome-lysosome fusion (Rap & CQN) impaired autophagic flux (**Figure 6D, E, Figure S10A**). Simvastatin or AY9944 treatment impaired autophagosome-lysosome fusion, leading to a reduction in autophagic flux comparable to Rap & CQN treatment (**Figure 6D, E, Figure S10A**). These results suggest that cholesterol depletion prevents autolysosome formation and possibly directs autophagosomes toward endosomal vesicles.

To determine if sterol depletion impacts autophagosome and late endosome fusion, we assessed colocalization of endosomal markers with LC3B (**Figure 6F, G, Figure S10B-D**). The late endosome marker Rab7 was highly colocalized with LC3B after simvastatin (∼40%) or AY9944 (∼60%) treatment (**Figure S10B, C**). Similarly, simvastatin and AY9944-treated cells exhibited ∼50% and ∼80% colocalization, respectively, between CD63 and LC3B expression compared to vehicle treatment, which showed <10% overlap in expression levels (**Figure 6F, G, Figure S10D**). We also observed significant overlap between Rab7, CD63, and LC3B expression (**Figure S10D**). Collectively, these findings suggest a mechanism whereby cholesterol deficiency prevents autophagosome-lysosome fusion, autophagic vesicles are directed toward late endosomes, and these amphisomes are then secreted from the cell as sEVs.

### Autophagy initiation is necessary for sEV secretion in cholesterol-depleted cells

Given the connection between autophagy, endosomal sorting, and sEV secretion, we predicted inhibition of autophagosome formation would prevent increased sEV release upon cholesterol depletion. ATG2 proteins function to expand and grow the developing and maturing autophagosome through lipid transfer to isolation membranes (Osawa et al., 2019; Tang et al., 2019a; Velikkakath et al., 2012). We utilized ATG2A/B knockout cells (HEK293T^ATG2A/B KO^), which exhibit limited autophagic activity due to the inability to transfer lipids to a growing phagophore (Tang et al., 2017; Tang *et al*., 2019a). To test if autophagosome maturation is necessary for cholesterol-deficient induction of sEV formation and release, we first analyzed CD63 and LC3B expression in HEK293T versus HEK293T^ATG2A/B KO^ cells. Both AY9944 and Rap & CQN increased CD63 expression concomitantly in HEK293T and HEK293T^ATG2A/B KO^ cells (**Figure 7A, B**), suggesting ATG2A/B KO does not prevent the accumulation of late endosomes following cholesterol depletion. Immunoblots of collected sEVs after pharmacological inhibition of cholesterol biosynthesis with simvastatin, AY9944, or U18666A showed increased LC3B expression relative to vehicle, suggesting increased secretion of LC3B-positive autophagic vesicles accompanies cholesterol depletion (**Figure 7D**). However, ATG2A/B KO prevented increased LC3B expression in response to AY9944 or Rap & CQN treatment (**Figure 7A, C**). Additionally, significant colocalization (∼80%) was observed between CD63 and LC3B after AY9944 or Rap & CQN treatments, an effect that was muted with ATG2A/B KO (**Figure 7A, E**). These data confirm ATG2A/B are required for proper autophagosome formation upon cholesterol depletion and ATG2A/B KO limits fusion events between late endosomes and autophagosomes. Lastly, we quantified the impact of ATG2A/B KO on cholesterol depletion-induced sEV secretion. While sEV secretion was increased after AY9944 or Rap & CQN treatment, this response was ablated in ATG2A/B KO cells (**Figure 7F, Figure S11**). Collectively, these findings support the concept that cholesterol deficiency drives sEV formation and release through recruitment of late endosome directed autophagic vesicles.

**Figure 7.**
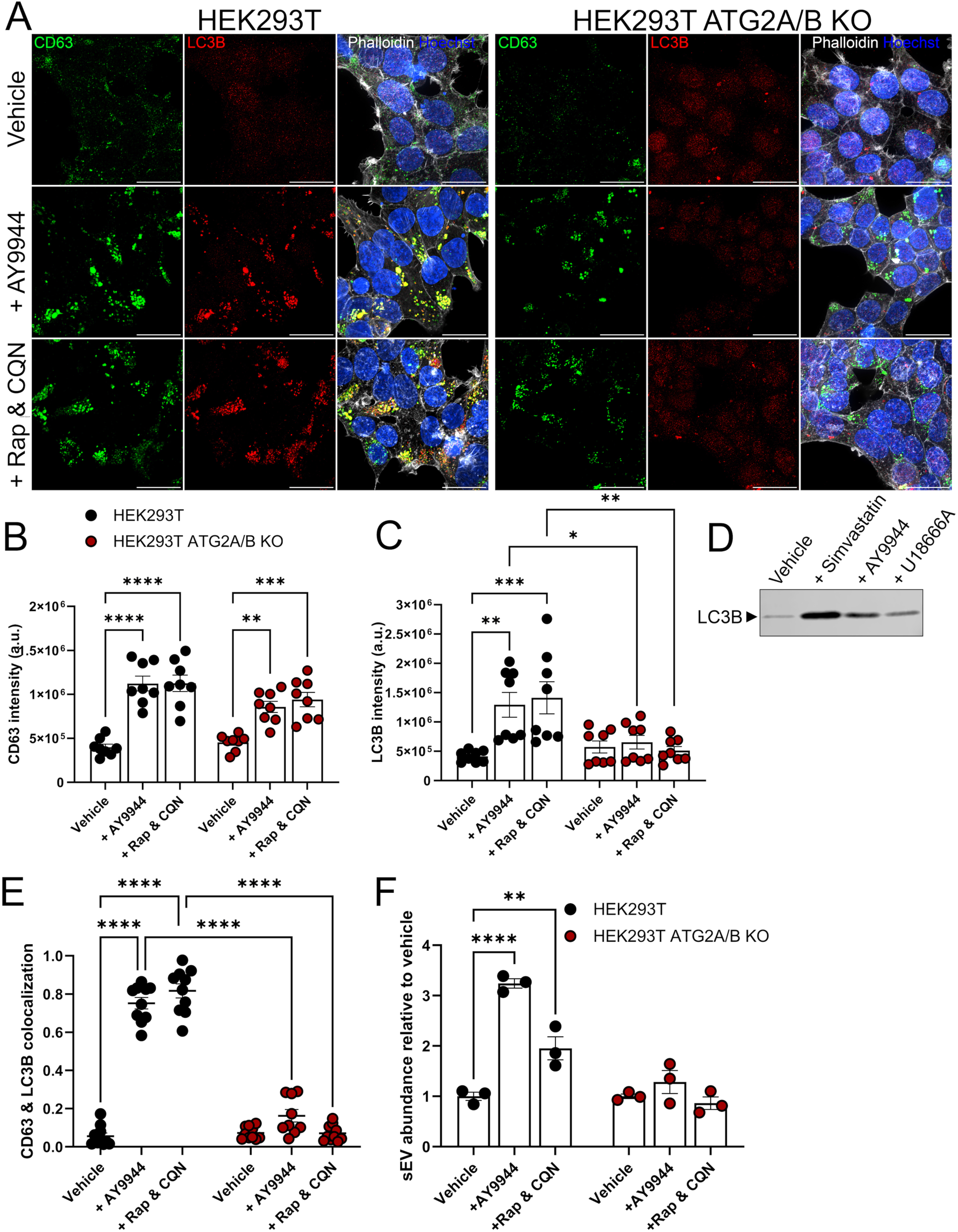
Increased sEV secretion in cholesterol-depleted conditions is due to autophagy-driven signaling. (**A**) Representative images for CD63 and LC3B expression in HEK293T or HEK293T ATG2A/B KO cells after vehicle, AY9944, or rapamycin and chloroquine (Rap & CQN) treatment. Scale bar, 25 µm. (**B**) Quantified CD63 expression in HEK293T and HEK293T ATG2A/B KO cells (mean ± SEM; n = 8 images analyzed from 2 independent experiments). Two-way ANOVA (Treatment effect: F2,42 = 47.99, p < 0.0001; Autophagy effect: F1,42 = 5.422, p ≤ 0.0248). Sidak’s multiple comparisons test (**p ≤ 0.005, ****p < 0.0001). (**C**) LC3B expression in HEK293T and HEK293T ATG2A/B KO cells (mean ± SEM; n = 8, 4 images analyzed from 2 independent experiments). Two-way ANOVA (Interaction effect: F2,42 = 6.126, p ≤ 0.0046; Treatment effect: F2,42 = 5.995, p ≤ 0.0051; Autophagy effect: F1,42 = 12.65, p ≤ 0.0009). Sidak’s multiple comparisons test (*p ≤ 0.05, **p ≤ 0.005, ***p ≤ 0.001). (**D**) Immunoblot for LC3B in isolated sEVs upon cholesterol depletion. (**E**) CD63 and LC3B colocalization in HEK293T and HEK293T ATG2A/B KO cells (mean ± SEM; n = 8 images analyzed from 2 independent experiments). Two-way ANOVA (Interaction effect: F2,53 = 128.9, p < 0.0001; Treatment effect: F2,53 = 157.0, p < 0.0001; Autophagy effect: F1,53 = 453.1, p < 0.0001). Sidak’s multiple comparisons test (****p < 0.0001). (**F**) Quantified sEV secretion in HEK293T or HEK293T ATG2A/B KO cells following drug treatments (mean ± SEM; n = 3 biological replicates from 3 independent experiments). One-way ANOVA (F2,6 = 56.17, p ≤ 0.0001), Dunnett’s multiple comparisons test; **p ≤ 0.01, ****p < 0.0001.

## DISCUSSION

Our results identify cholesterol homeostasis as a critical regulator of sEV biology. Through use of pharmacological inhibitors and genetic models of cholesterol biosynthesis disorders, cholesterol disruption increases sEV release through autophagy-directed vesicle buildup and release. Increased biogenesis of sEVs in cholesterol-deficient conditions negatively correlated to cholesterol levels across cell models (**Figure 1H, Figure S4D, E**). Previous reports have suggested cholesterol metabolism to be an essential feature of sEV formation and release (Kulshreshtha *et al*., 2019; Sekhavati *et al*., 2023). Our data complements recent studies demonstrating statins enhance sEV secretion (Martin et al., 2023) while excessive cholesterol (hypercholesterolemia) impairs sEV release (Abdullah et al., 2021). Distinct pathways have been introduced within the context of sEV biogenesis, including those necessitating endosomal sorting complexes required for transport (ESCRT) machinery and those that may be produced independently of ESCRT proteins, including tetraspanin- and flotillin-dependent pathways (Kenific et al., 2021; Larios et al., 2020; van Niel *et al*., 2018). Although cholesterol appears to play an essential role in the overall function of flotillin proteins (Kwiatkowska et al., 2020), biogenesis of ESCRT-derived sEVs do not require cholesterol (Wei et al., 2021). Further, sEV-associated tetraspanin proteins may partly depend on cholesterol to fulfill certain functions. For example, CD63, a component of the flotillin-dependent sEV biogenesis pathway, is involved in sorting cholesterol into ILVs (Palmulli *et al*., 2024). CD81, involved in tetraspanin-dependent sEV formation, has a protein structure that contains a cholesterol binding pocket (Zimmerman et al., 2016). These features may partially explain why CD63 and CD81 surface markers are slightly reduced in sEVs produced from cells treated with AY9944 (**Figure 2B, D**). However, CD9 clustering in late endosomes is sufficient for inducing membrane bending essential in forming ILV invaginations (Umeda et al., 2020), a function that corresponds to unchanged surface marker levels in our study (**Figure 1F**, **Figure 2B, D**).

While sEV release is increased after disrupted cholesterol biosynthesis, our data suggest the secreted products are structurally altered relative to control conditions (**Figure 3**). Physically dysmorphic sEVs generated after cholesterol starvation are reminiscent of misshapen clathrin-coated pit structures previously observed under cholesterol-depleted condition (Anderson *et al*., 2021). Occurrence of malformed vesicles in both of these pathways is presumably a consequence of reduced membrane-associated cholesterol. Cholesterol distribution during sEV biogenesis is not fully understood, but most of the current evidence is derived from studies of the cholesterol storage disorder, Niemann-Pick disease, Type C1. Endosomal cholesterol is present in two varieties: plasma membrane derivatives and sterols collected from low density lipoproteins (LDL). Cholesteryl ester-rich LDL is taken up through receptor-mediated endocytosis before it is hydrolyzed and temporarily stored as free cholesterol in lysobisphosphatidic acid-containing late endosomes (Kobayashi et al., 1999). Accumulation of late endosomal free cholesterol does not affect the abundance of lipid rafts (Sobo et al., 2007); instead, free cholesterol is transported from late endosomes/lysosomes through vesicles or transport proteins to organelles, most significantly the ER (Luo et al., 2017). In cholesterol deficient conditions, limited plasma membrane- and LDL-supplied cholesterol likely leads to structurally challenged sEV formations, which may ultimately impact surface marker expression and internalization (**Figure 2**, **Figure 3**, **Figure 4**).

In our study, we observed alterations of sEV surface marker expression after cholesterol biosynthesis disruption (**Figure 2, Table S1**). While protein expression on EVs may define their origin as ectosomes or exosomes (Mathieu et al., 2021), size-discriminating NanoFACS findings (**Figure 2F, Figure S7**) and sEV expression profiles demonstrate the majority under study were formed intracellularly. ROR1 (receptor tyrosine kinase-like receptor 1) expression was significantly increased in all cholesterol-targeted treatment groups (**Figure 2C, D, E, Table S1**). As ROR1 triggers non-canonical Wnt signaling by binding Wnt5a (Fukuda et al., 2008), its increased packaging within sEVs could be an attempted compensatory mechanism to normalize impaired Wnt signaling accompanying cholesterol biosynthesis disruption (Francis et al., 2016; Sengupta et al., 2024). In a similar vein, increased packaging of CD29, an integrin important for mediating cell adhesion properties, could be related to elevated internalization of sEVs released from cholesterol deficient cells (**Figure 4**). Importantly, the modifications to sEV surface marker and sterol profiles seen here, may further coincide with changes in transported nucleic acids (Lee et al., 2024).

Despite deleterious effects on clathrin-mediated endocytosis (Anderson *et al*., 2021), cholesterol biosynthesis impairments seem to stimulate an increase in endosome formation (**Figure 5, Figure S9**). We surmise this results from a roadblock in endosomal sorting pathways, perhaps associated with defects in docking or fusion mechanisms (Luzio et al., 2007). Notably, Rab7 plays important roles in late endosome-lysosome fusion, lysosome formation, and lysosome preservation (Bucci et al., 2000; Zhang et al., 2009). As such, Rab7 should thus not be a definitive marker for late endosomes. The spatial differences between lysosomes relative to Rab7-positive vesicles, together with the colocalization of Rab7 with CD63 (**Figure S9B, Figure S10B, D**), suggest Rab7 is labeling late endosomes in cholesterol-depleted conditions. Alongside its connection to lysosomes, Rab7 mediates late endosome-autophagosome fusion events (Jäger et al., 2004), an effect we observed with cholesterol depletion (**Figure S10B, C**). While mature autophagosomes typically fuse with lysosomes, we demonstrate autophagosomes are redirected to late endosomes in cholesterol deficient cells (**Figure 6, Figure S10**). While suppression of autophagic flux could be linked to defects in the functions of soluble N-ethylmaleimide-sensitive factor activating protein receptor (SNARE) complexes, Rab proteins, or phosphoinositide 3-kinase (PI3K) activity (Hikita et al., 2018), additional studies are needed to uncover the precise mechanisms involved.

A critical finding of our study is the connection between autophagosome maturation and sEV secretion during cholesterol depletion (**Figure 7**). These results are significant for several reasons. First, our data link cholesterol homeostasis and autophagy which has not previously been thoroughly described (Russell et al., 2014). Second, our work goes against the assumption that as a major lipid component of sEVs, cholesterol is required for sEV release. Lastly, our data suggest a novel cellular mechanism by which cholesterol content regulates sEV secretion. In its most simplistic form, the initiation of autophagosome formation involves the transport of an isolation membrane to a membrane-rich source (like the ER) and the shuttling of lipids to a growing phagophore (Yamamoto et al., 2023). Recent work has linked autophagosome formation to sEV secretion (Kim et al., 2021; Solvik et al., 2022; Wei et al., 2022). In our study, we prevented phagophore expansion by deletion of ATG2A and ATG2B (Tang *et al*., 2019a; Tang et al., 2019b), inhibiting late endosome-autophagosome fusion during cholesterol depletion (**Figure 7A, E**). While deletion of ATG2A and ATG2B prevented autophagosome accumulation (**Figure 7C**), it did not correct the increase in CD63-positive vesicles that accompanied cholesterol biosynthesis inhibition (**Figure 7B**). This finding suggests that the accumulation of late endosomes, in response to cholesterol disruption or treatment with rapamycin and chloroquine, is not directly tied to autophagosome formation. However, sEVs secreted from cholesterol-depleted cells expressed LC3B (**Figure 7D**), an effect also observed in autophagy-stimulated cells (Solvik *et al*., 2022). Prevention of autophagosome elongation by ATG2A/B deletion impaired sEV secretion due to cholesterol biosynthesis inhibition (**Figure 7F**), implying that autophagic vesicles are an inseparable component of sEV release during cholesterol depletion.

Many disease states are associated with cholesterol imbalance (Dai *et al*., 2021). Here, we report fibroblasts derived from subjects with two rare disorders of cholesterol biosynthesis, SLOS and lathosterolosis, release increased numbers of sEVs relative to unaffected controls (**Figure 1C**). Previous work on SLOS fibroblasts identified multilamellar structures (Wassif et al., 2002) which resemble formations observed in our study (**Figure 5F**). Our results suggest these structures are likely a consequence of autophagic vesicle redirection. This idea concurs with identified phagocytic structures in the retinal pigment epithelium of AY9944-treated rats (Fliesler et al., 2018), as well as the accumulation of immature autophagosomes in induced pluripotent stem cell (iPSC)-derived SLOS retinal pigment epithelium cells (Ramachandra Rao et al., 2018). The mechanisms linking cholesterol homeostasis, autophagy, and sEV secretion are also likely translatable to other common diseases. For example, Alzheimer’s disease is associated with cholesterol disequilibrium (Xue-shan *et al*., 2016), autophagy abnormalities (Nixon and Rubinsztein, 2024), and distinguished sEV dynamics (Liang et al., 2023). Translation of our findings to this and other cholesterol-impacted diseases should be explored.

In summary, we have uncovered an interplay between cholesterol metabolism and sEV release, highlighted structural and phenotypic alterations in sEV biology as a consequence of cholesterol depletion, characterized internalization of cholesterol-depleted sEVs, and defined the intracellular mechanism by which autophagy drives sEV release in cholesterol-depleted cells. While the downstream biological significance of cholesterol impacts on sEV biology and disease implications require further elucidation, our study provides a critical mechanistic-driven framework from which these ideas can be further investigated.

## ACKNOWLEDGEMENTS

This study was supported by the National Institutes of Health under Award Numbers R01DE032712, U54GM128729, P20GM103620 and P20GM103548. We would like to thank Hong-Gang Wang (Pennsylvania State University) for the kind gift of ATG2A/B knockout cells. We thank the Mayo Microscopy and Cell Analysis Core for experimental and technical support. We would like to thank the University of South Dakota Center for Brain and Behavior Research for supporting this project. We also thank Kelly Graber within the Histology and Imaging Core at Sanford Research, as well as Jared Wollman within the Flow Cytometry Core at Sanford Research, for their technical expertise. Illustrations presented in figures were created with BioRender. Any opinions, findings and conclusions expressed in this material are those of the author(s) and do not necessarily reflect the views of the National Institutes of Health.

## AUTHOR CONTRIBUTIONS

Conceptualization, J.D.W.Y. and K.R.F.; Methodology, J.D.W.Y., S.S., A.L.W. and M.M.; Investigation, J.D.W.Y., S.S., A.L.W., and M.M.; Writing – original draft, J.D.W.Y.; Writing – review & editing, J.D.W.Y., S.S., A.L.W., M.M., T.K.M., P.D.V., and K.R.F.; Funding acquisition, P.D.V. and K.R.F.; Supervision, T.K.M., P.D.V., and K.R.F.

## DECLARATION OF INTERESTS

The authors declare no competing or financial interests.

## MATERIALS AND METHODS

### Cell culture

Fibroblasts, collected from skin punch biopsies from control, Smith-Lemli-Opitz syndrome (SLOS), and lathosterolosis (LATH) subjects (kind gift from Dr. Forbes Porter, NICHD), were cultured in DMEM (GIBCO, 4.5 g/L glucose, 110 mg/L pyruvate) mixed with 15% (v/v) fetal bovine serum (FBS; Hyclone, SH3039603), and 1,000 U/mL penicillin/streptomycin (Life Technologies, 15140122). To force cells to utilize endogenous cholesterol synthesis mechanisms and prevent lipoprotein-mediated internalization from the culture medium, the medium containing FBS was replaced with lipoprotein-deficient serum (LPDS) 48 h after seeding. Fibroblasts were rinsed twice in PBS and grown for 7 days in 7.5% (v/v) LPDS to induce biochemical profiles. SLOS and LATH fibroblast cell lines have been described previously (Krakowiak *et al*., 2003; Wassif *et al*., 1998). Immortalized human embryonic kidney cells (HEK293T) and keratinocytes (HaCaT) were cultured in DMEM (GIBCO, 4.5 g/L glucose, 110 mg/L pyruvate) combined with 10% (v/v) FBS (Hyclone), and 1,000 U/mL penicillin/streptomycin. Similarly, immortalized human microglia (HMC3) were grown in EMEM (ATCC, 30-2003) containing 10% (v/v) FBS and 1% (v/v) penicillin/streptomycin. Cells were allowed 24 h to adhere to culture treated dishes, after which time media was replaced with 10% (v/v) LPDS conditions to prompt cholesterol biosynthetic processes. Inhibition of cholesterol biosynthesis included cells being rinsed in phosphate buffer solution (PBS) and cultured for 48 h in LPDS followed by 24 h or 48 h treatment with small molecule inhibitors AY9944 (2.5 μM, Cayman Chemical; DHCR7 inhibitor), simvastatin (1 μM, Cayman Chemical; HMG-CoA Reductase inhibitor), U18666A (20 nM, Cayman Chemical; DHCR24 inhibitor), Avasimibe (5 μM, Cayman Chemical; ACAT inhibitor), or GW4869 (2.5 μM, Cayman Chemical; neutral sphingomyelinase inhibitor).

### Generation of overexpression and knockout cell lines

Plasmids housing genes for proteins involved in sEV biogenesis or mature autophagosome formation fused to fluorescent reporters, including mEmerald-CD9 (a gift from Michael Davidson, Addgene plasmid # 54029), eGFP-CD63 (a gift from Paul Luzio, Addgene plasmid # 62964), and mCherry-LC3B (a gift from David Rubinsztein, Addgene plasmid # 40827) were isolated and purified (GeneJET Plasmid Maxiprep Kit, Cat. # K0491). These plasmids possess similar backbones with a CMV promoter, kanamycin or ampicillin bacterial resistance, and neomycin selection markers. Additionally, an autophagic flux reporter line was generated utilizing a GFP-LC3-RFP-LC3ΔG plasmid (a gift from Noboru Mizushima, Addgene plasmid # 84572), which has been previously described (Kaizuka *et al*., 2016). To produce stably transfected cells, HEK293T cells were seeded onto 0.1% gelatin-coated six well plates at a density of 300,000 cells/well. After 24 h, cells were transfected using Lipofectamine 2000 (ThermoFisher Scientific, Cat. # 11668019) per manufacturer’s instructions. In short, 250 μL Opti-Mem (ThermoFisher Scientific, Cat. # 31985062) was combined with 4 µg of plasmid DNA in one solution and with 10 μL of Lipofectamine 2000 in a second solution. Both mixtures were allowed to incubate for 5 min at room temperature. Solutions of DNA/Opti-MEM and Lipofectamine/Opti-MEM were then combined with gentle mixing and allowed to incubate at room temperature for 20 minutes. Fresh media was added to cells followed by the addition of 500 μL of the transfection mixture in a dropwise fashion. Cells were allowed to incubate at 37°C for 6 h after which the media was changed. To produce stable cell lines, neomycin or puromycin was added to the media at a concentration of 1.75 mg/mL or 2.5 µg/ml for 7 days. To maintain the stable transfection in culture, neomycin or puromycin was periodically reintroduced to cells that have undergone several passages. To explore how early autophagy formation effects extracellular vesicle formation in the context of cholesterol starvation, we utilized ATG2A/B knockout cells (HEK293T^ATG2A/B KO^), a kind gift from Hong-Gang Wang (Pennsylvania State University), which have been previously described (Tang *et al*., 2017; Tang *et al*., 2019a).

### Preparation of lipoprotein-deficient serum (LPDS)

As cells can internalize cholesterol from the environment, we produced fetal bovine serum (FBS) lacking sterols, triglycerides, and other neutral lipids to promote de novo sterol synthesis based on techniques described elsewhere (Cham and Knowles, 1976). In brief, oxidation caused by trace peroxides was prevented by adding 0.1 mg ethylenediamine tetraacetate (EDTA) per 50 mL FBS. Serum was then combined with an organic phase (3:2 ratio diisopropyl ether:n-butanol) in a 1:2 mixture with vigorous stirring for 1 h protected from light. The bulk organic phase was discarded and the remaining mixture was centrifuged at 4°C for 15 min at 2200 rpm after which the aqueous layer was collected, filtered, and lyophilized. The lyophilized product was then resuspended in molecular grade H_2_O and insulin-transferrin-selenium (ITS-G) supplement (Gibco, 41400045) was added. The solution was filter-sterilized and stored at 20°C. We used GC-MS to validate cholesterol levels that were nearly undetectable in the produced lipoprotein-deficient serum (LPDS).

### Gas chromatography-mass spectrometry (GC-MS)

Cell pellets were collected and flash frozen on dry ice before being reconstituted in 1 mL of H_2_O and lysed by successive freeze/thaw cycles between dry ice and a 50°C bead bath. Protein quantification (Micro BCA Protein Assay Kit, ThermoFisher, 23235) was performed using 50 μL from each sample, and 1 mL of saponification buffer (7% KOH, 92% ethanol, 10 μg/mL coprostan-3-ol) was added to the remaining cell lysate (∼900 μL). Saponification took place at 60°C for 1 h after which an additional 1 mL of water was added to each sample. Separation of the aqueous phase was initiated with 3 mL ethyl acetate and vortexing before centrifugation for 5 min at 2200 rpm. After organic phase extraction in 2 mL water, each sample was concentrated to dryness by applying heat (50°C) under constant nitrogen flow. Residual sterols were then dissolved in 50 μL pyridine (CHROMASOLV Plus, Sigma) and derivatized in 50 μL N,O-bis(trimethylsilyl) trifluoroacetamide with 1% trimethylchlorosilane (BSTFA + 1% TMCS, ThermoFisher TS-38831) at 50°C for 1 h. Sterol analysis was performed by GC-MS. An autosampler injected 1 μL of each derivatized sterol sample into an Agilent 7890 GC using a split injection port (4 mm ID x 78.5 mm quartz wool liner, Restek 23309) connected to a 0.18 mm ID x 20 m 1,4-bis(dimethylsiloxy)phenylene dimethyl polysiloxane column (Restek, 43602). The carrier gas was helium which flowed at a linear rate of 46.9 cm/ sec. The method produced for sample analysis progressed as follows: 0.5 min at 170°C, oven temperature was then raised to 250°C at 18°C/min, followed by an increase to 280°C at 3°C/min, and finally to 320°C at 20°C/min where the temperature was held for 7 min. A mass spectrometer (Agilent 5977B) was operated in electron impact mode at 70 eV at an ion source temperature of 275°C.

### Sterol quantification by GC-MS

MassHunter software was used for GC-MS analysis. TMS-derived ethers of natural sterols were identified through comparison of commercially available standards for cholesterol, 7-dehydrocholesterol (7-DHC), lathosterol, and desmosterol (Avanti Polar Lipids, Inc.), but also through comparison to MS spectra available through the National Institute of Standards and Technologies Standard Reference Database when available. The sterol 8-dehydrocholesterol (8-DHC) was identified as an isomer of 7-DHC and compared to SLOS fibroblasts. Zymostenol was recognized from spectra of Conradi-Hünermann-Happle syndrome (CDPXD2) fibroblasts. Representative fragmentation patterns from spectra are available upon request. Normalization of sterol abundance was performed using both the internal standard (coprostanol) and protein concentration (Micro BCA Protein Assay Kit, ThermoFisher, 23235), and these data were analyzed as relative to control samples using GraphPad Prism software.

### Small extracellular vesicle (sEV) isolation

Cells were grown in 10 cm dishes where 8 mL of media that was depleted of extracellular vesicles (centrifugation for 16 h at 110,000 x g) (**Figure S1A**) was provided 48 h prior to sEV isolation, which was performed using differential centrifugation that has been similarly described (Wang et al., 2021). Media from each 10 cm dish was collected and spun at 2,000 x g for 20 min to remove cell debris and large particles, including most apoptotic bodies. The supernatant from two 10 cm dish samples were combined and centrifuged at 10,000 x g for 30 min (4°C) to pellet moderately sized particles, including microvesicles. The solution above the microscopic pellet was carefully removed, placed into pollyallomer tubes (Thermo Scientific, Cat. # 03126) and spun using an ultracentrifuge (Thermo Scientific Sorvall WX80 Ultracentrifuge) set at 110,000 x g for 2 h (4°C). The supernatant was carefully aspirated and the pelleted sEVs were washed with PBS on ice for 10 min. The sample was centrifuged again (110,000 x g, 2 h, 4°C) after which the supernatant was carefully aspirated and isolated sEVs were incubated in 300 μL PBS at room temperature for 15 min, after which they were used immediately or stored in 50 μL aliquots at -80°C.

### Quantification of sEVs

Fresh or frozen aliquots of sEVs were diluted in PBS (typically 1:10, but higher dilutions when particle concentrations were high) and flowed at a steady rate (NanoSight syringe pump) through a NanoSight NS300 (Malvern Panalytical) instrument. Particles from each sample were visualized and quantified as five technical replicates using Nanoparticle Tracking Analysis (NTA) software. Exported results were copied into GraphPad Prism for statistical comparisons. NanoSight equipment was thoroughly cleaned with nanopure H_2_O between samples and 10% ethanol at the end of each analysis session. The particle size distributions were calibrated using calibration standards. Concentrations of sEVs were normalized by cell count data collected using cell counting chamber slides (ThermoFisher, C10228) and Countess II Automated Cell Counter (ThermoFisher). Cell viability was measured using 0.4% trypan blue staining with the Countess Cell Counter system.

### Exosome population assay and flow cytometry

sEV marker changes associated with treatments were assessed using MACSPlex human exosome kit (Miltenyi Biotec) per manufacturer’s instructions. In brief, 2.5 × 10^8^ sEVs (quantified by NanoSight analyses described above) were diluted in MACSPlex buffer to a final volume of 120 μL. A sample without sEVs and only buffer was used as a blank control. Exosome capture beads were vortexed for 30 s prior to 15 μL being added to each sample (or blank) tube. Tubes were incubated overnight in a thermomixer set to 450 rpm at 20°C protected from light. The next day, 500 μL of buffer was added to each tube followed by centrifugation at 3000 x g for 5 min. Next, 500 μL of the supernatant was carefully removed and 15 μL from a cocktail containing 5 μL of each MACSPLEX detection reagent (CD9, CD63, and CD81) per sample was mixed into each tube. Protected from light, samples were incubated at room temperature and mixed at 450 rpm in a thermomixer for 1 h. Following the addition of 500 μL wash buffer, tubes were centrifuged again at 3000 x g for 5 min, and 500 μL of the supernatant was again carefully aspirated. The final steps included a secondary wash with 500 μL buffer, 15 min incubation in a thermocycler set to room temperature and 450 rpm, 5 min centrifugation at 3000 x g, and aspiration of 500 μL, which left a final product that was diluted with 50 μL producing a final volume of 200 μL ready for measurement with a flow cytometer. A Fortessa flow cytometer (BD FACS LSRFortessa SORP) was primed to remove bubbles and cleaned with FACS clean, FACS rinse, and ddH_2_O prior to use. Tracking beads were used to align lasers, set PMT voltages, set delays, and compare against previous baseline measurements to insure system consistency. MACSPlex setup beads were used to appropriately gate bead populations and discern single events from debris or events that include clumping of multiple beads. For each sample, events were recorded for FITC vs PE fluorescence. The median fluorescence intensity of each sEV population of the blank control sample was subtracted from that of each experimental sample to produce a background-subtracted value that was used for analyses. While the MACSPlex human exosome kit allows detection of up to 37 distinct surface epitopes on sEVs, we report only populations with signal intensities over 400; however, we did include REA and mIgG1 controls to show that non-specific binding was minimal in our assays. Data was analyzed using GraphPad Prism and FlowJo software.

### NanoFACS analysis of small and medium sized EVs

Samples were tested while blinded to phenotype. They were first centrifuged at 2500xg x 15 min per standard ISEV recommendations (Welsh et al., 2020). The supernatant was then diluted 1:200 into 200 nm bead buffer (Fluoresbrite® YG Microspheres (200 nm), Polysciences) created by FACS sorting 200 nm beads into 0.1 µm filtered PBS. Polystyrene 200 nm bead buffer provided both a relative internal size estimate and a means to standardize the volume of sample tested (e.g. 2000 beads equaled 1 µL of starting sample tested on our FACSAria Fusion (BD Biosciences). NanoFACS data was gated relative to the internal 200 nm as “small: < 200 nm” and “large: 200-900 nm”. Briefly, all experiments were performed in triplicate as previously described by our group (Clemente et al., 2020; Kim et al., 2022; Kuhn et al., 2022; Morgan, 2018). NanoFACS relies on either relative fluorescence for sizing EVs (Welsh *et al*., 2020) or side light scatter (SSC-H), which is highly dependent on the diameter of the particle and wavelength of the laser with some contribution based on refractive index. Notably, small EVs have a refractive index more like a virus (1.58) than a cell (1.38) compared with polystyrene bead controls (1.6). Larger EVs have RI closer to a cell leading to an underestimate of their true size relative to bead (∼2-4 fold). Machine settings were standardized for experiments in triplicate using commercially available polystyrene bead standards ranging in size from 100 nm to 900 nm, Megamix-Plus FSC and -Plus SSC, BioCytex). Fluidic and instrument settings were designed to minimize background noise while maximizing the detection of submicron sized EVs. Imaging was performed on a FACSAria Fusion (BD Biosciences) with a 70 µm nozzle and 0.1 µm filtered PBS sheath fluid at a pressure of 70 psi. The FacsAria Fusion employs a 50-mW 488-nm blue laser and SSC photomultiplier (PMT) with 488/10 BP filter, and a 505 splitter with a 525/50 BP filter for the detection of GFP labeled beads. SSC-H voltage was uniformly adjusted for all experiments so 200 nm beads were at SSC 10^4^ and 900 nm beads remained on scale. All flow machine buffers were filtered with a ZenPure PureFlow Mini Capsule PES 0.1 µm filter. Coincident EV events per nanoliter droplet were controlled by using a threshold rate of < 40,000 events per second and electronic abort rate less than 5%. An essential criterion for reproducibility was setting uniform gates in advance of batch analysis with positive and negative control samples [1.6 µl of Triton X-100 (10% aqueous solution, Sigma-Aldrich) was added to the remaining sample (∼160 µL) for a final detergent concentration of 0.1%, gently mixed by pipetting, and run again using the same settings as DETERGENT negative controls for each sample]. All samples used the same pre-set gates distinguishing < 200 nm vs 200-900 nm. Data were collected based on uniform 200 nm bead counts spiked into each sample to standardized plasma volumes being tested (data reported as counts/µL of plasma). The mean of triplicate experiments for each sample was adjusted by corresponding bead counts (2000 beads counted/µL) were employed for statistical analysis. Results were reported as the mean +/-95% confidence intervals analyzed by ANOVA with *post hoc* verification compared with vehicle and exposure.

### Immunocytochemistry and confocal microscopy

Cells were grown on 12 mm glass coverslips coated with 0.25% fibronectin. Before antibody staining, cells were fixed in 4% paraformaldehyde (PFA; Electron Microscopy Sciences, 15714) at room temperature for 20 min. Next, cells were permeabilized with 0.2% Triton X-100 (Sigma-Aldrich, 93443) for 20 min and then incubated in blocking buffer (5% donkey or goat serum, Jackson ImmunoResearch Laboratories, 017-000-121 or 005-000-121; 0.1% Triton X-100) for 1 h at room temperature. The primary antibodies used for these studies were diluted in blocking buffer and included rabbit anti-Rab5 (Cell Signaling Technologies, 3547, 1:200), rabbit anti-Rab7 (Cell Signaling Technologies, 9367, 1:200), rabbit anti-LC3B (Cell Signaling Technologies, 3868, 1:200), rabbit anti-LAMP1 (Novus, 19294, 1:1000), and mouse anti-CD63 (Novus, 77913, 1:500). Primary antibodies were visualized with Alexa Fluor-conjugated secondary antibodies (Life Technologies, A11001, A21437 and A31570; 1:500). Hoechst 33342 nuclear counterstain (Invitrogen, H3570, 1:10,000), and phalloidin conjugated to a far-red Alexa 647 dye (ThermoFisher, A22287, 1:400) were incubated with the secondary antibodies where indicated. Images were acquired using a Nikon A1R resonant scanning multispectral confocal microscope (Nikon Instruments, Inc. Melville, NY) or Nikon CSU-W1 Spinning Disk Super Resolution by Optical Pixel Reassignment (SoRa) system, equipped with NIS-Elements analysis software (Nikon).

### sEV internalization assays

HEK293T^mEmerald-CD9^ and HEK293T^eGFP-CD63^ cells overexpressing common sEV markers were treated with cholesterol biosynthesis inhibitors in LPDS for 48 h before sEVs were isolated from the media and quantified (described above). HEK293T cells were plated onto fibronectin-coated (0.25%) coverslips (20,000 cells/coverslip) and allowed 24 h to attach. Next, cells were washed gently with PBS and media was replaced with DMEM containing 1×10^8^ or 2.5×10^8^ sEVs from vehicle-, simvastatin-, or AY9944-treated cells overexpressing sEV markers (HEK293T^mEmerald-CD9^ or HEK293T^eGFP-CD63^). These values were chosen from sEV dose response experiments (**Figure S8**). Cells were allowed to incubate (37 °C, 5% CO_2_) for 12 h, determined based on previously described methods (Jurgielewicz et al., 2020). After incubation, cells were washed thoroughly with PBS and stained with 594-conjugated concanavalin A (ConA; Biotium, 29017, 100 µg/mL) and Hoechst (1:10,000) diluted in HBSS for 30 min. Cells were then washed twice with HBSS and fixed with 4% PFA for 15 min at room temperature. Upon completion of fixation, coverslips were washed three times and mounted (Vector Lab., Vectashield mounting medium, H-1700-10) for imaging. Cells were imaged on Nikon CSU-W1 SoRa microscope with a 60x objective and 2.8x magnification. Z-stack images were collected using serial optical dissections set at a 200 nm thickness. 3D renderings were generated using NIS Elements AR. Internalized particle detection was quantified using ImageJ particle analysis where a set detection threshold was utilized for each experiment.

### Transmission electron microscopy (TEM)

Frozen aliquots of sEVs isolated from HEK293T cells treated with vehicle, simvastatin, or AY9944 were prepared for transmission electron microscopy by applying 3 µl to formvar/carbon coated grids (EMS, Hatfield, PA) after which they were allowed to adhere for several minutes. Following two brief rinses in H_2_O, 1% phosphotungstic acid was applied to grids and allowed to dry prior to imaging. HEK293T cells pellets were collected and fixed in Trump’s Fixative, which was prepared by dissolving monobasic sodium phosphate (Na2HPO4H2O) and sodium hydroxide (NaOH) in distilled water before adding glutaraldehyde, formaldehyde, and hydrochloric acid (HCl) to bring the pH to 7.2. The final product is a 4% formaldehyde, 1% glutaraldehyde in 0.1 M phosphate buffer. After fixation, cell pellets were suspended in 2% low melt agar for further processing. Briefly, cells were stained with osmium tetroxide, washed, stained with 2% uranyl acetate, washed and dehydrated through a graded ethanol series followed by embedding in Embed 812 resin (EMS). After a 24 h polymerization at 60°C, 0.1 μM ultrathin sections were prepared and post-stained with lead citrate. All micrographs were acquired using a JEOL 1400 Plus TEM (JEOL USA, Peabody, MA) operating at 80 kV and equipped with a Gatan Orius camera (Gatan, Inc., Warrendale, PA).

### Western blot analyses

Previously quantified EV samples (see above section) were lysed in 2x Laemmli sample loading buffer (BioRad, 1610737) with 4% β-mercaptoethanol at 95°C for 10 minutes. A total of 1.0×10^8^ (CD9) or 7.5×10^8^ (CD29, ROR1, LC3B) sEVs were loaded per lane of 4-20% Mini-PROTEAN precast protein gels (BioRad, 4561094). Gels were blotted onto PVDF membranes (Fisher Scientific, IPVH00010) and membranes were blocked in 5% milk in TBS-T (Tris-buffered saline-Tween 20) for 1 h. Primary antibody incubation was carried out in 1% blocking solution overnight at 4°C (rabbit anti-CD9, Cell Signaling Technologies, D3H3P, 1:1000; mouse anti-CD29, BD Biosciences, 610467, 1:2500; goat anti-ROR1, Novus, AF2000, 1:500; rabbit anti-LC3B, Cell Signaling Technologies, 3868, 1:1000). Secondary antibody incubation was performed for 1 h at room temperature (Jackson ImmunoResearch, 711-035-151, 711-035-152,705-035-003). Blots were developed and exposed on a Li-Cor Odyssey FC using Immobilon Classico Western HRP substrate (Millipore Sigma, WBLUC0100). Precision Plus Protein Kaleidoscope pre-stained protein ladder (BioRad, 1610375) was used for determining molecular weight.

### Statistical analysis

All statistical analysis was performed using GraphPad Prism 8.0.2 (GraphPad Software, Inc., CA, US). Homogeneity of variances was tested by the Brown-Forsythe test. In cases of equal variances, data was analyzed by one-way ANOVA and *post hoc* Dunnett t test for multiple comparisons relative to vehicle control group. In cases of unequal variances, Welch ANOVA and *post hoc* Dunnett’s T3 test was performed. Correlations were performed using simple linear regressions where a significant slope value is determined to be different from zero. For significant correlations, the R-squared and p-values are reported. Colocalization analyses were performed using a Fiji (ImageJ 1.54f) plugin called just another colocalization plugin (JaCoP), where Mander’s coefficient values (the proportion of overlap between channels) were determined based on a set threshold that remained unchanged for each experiment. P-values of < 0.05 were accepted as significant. Statistical details for each experiment can be found in the figure legends.

## Notes

### Competing Interest Statement

The authors have declared no competing interest.

