## Supplemental Info and Figures for "Cholesterol deficiency directs autophagy-dependent secretion of extracellular vesicles"

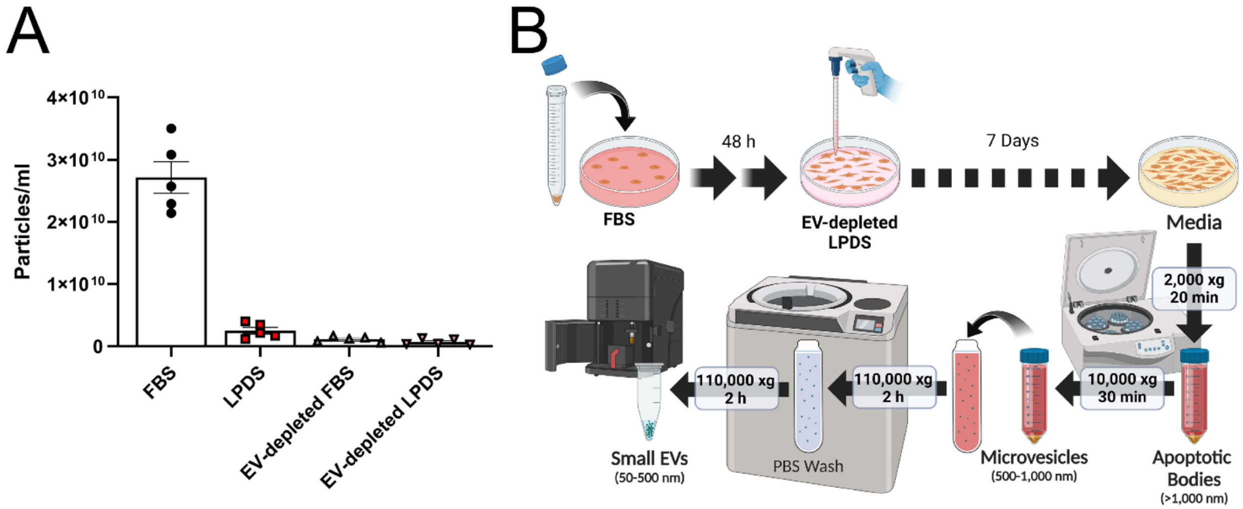


**Figure S1. Validation of sEV depletion of serum and sEV isolation protocol. Related to Figure 1.**

(A) Nanoparticle tracking analysis (NTA) of sEVs in FBS, lipoprotein-deficient serum (LPDS), and EV-depleted (ultracentrifugation at 110,000xg for 16 h) conditions (mean ± SEM; n = 5).

(B) Experimental plan for fibroblast experiments where FBS media was replaced after 48 h with EV-depleted LPDS media. After 7 days, media was collected, sEVs were isolated with differential centrifugation, and quantified with NTA.

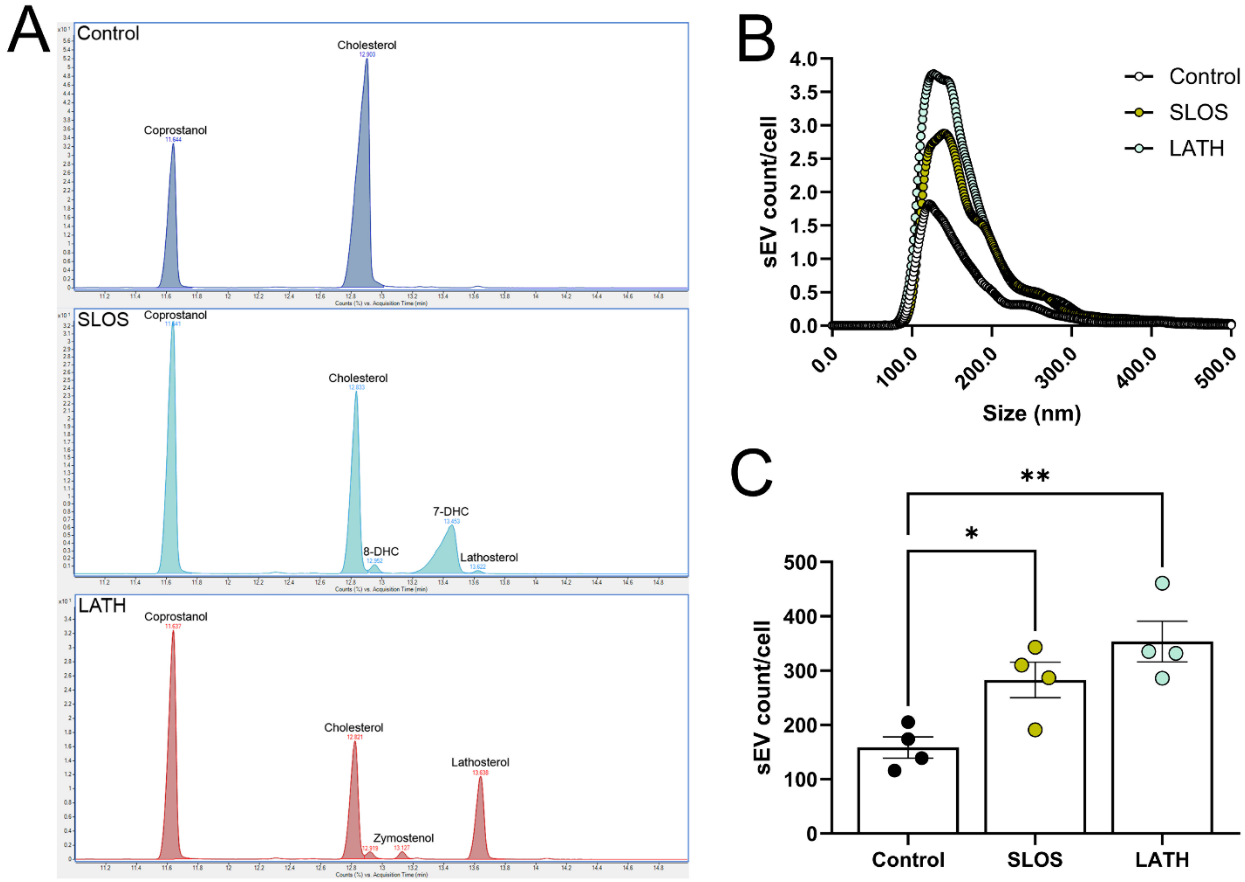


**Figure S2. Patient-derived fibroblasts exhibiting mutations in cholesterol biosynthetic genes exhibit impaired cholesterol production and increased sEV secretion. Related to Figure 1.**

(A) Chromatograms from GC-MS analyses of fibroblasts reveal disease-associated biochemical changes in SLOS and LATH patient-derived samples in the form of reduced cholesterol expression and accumulation of sterol intermediates. Coprostanol is used as an internal standard.

(B) NTA of sEVs shows a similar size range (~80 – 300 nm) for control, SLOS, and LATH fibroblasts (mean; n = 4 biological replicates from 4 independent experiments).

(C) SLOS and LATH fibroblasts secrete more sEVs compared to unaffected controls (mean ± SEM; n = 4 biological replicates from 4 independent experiments). sEVs are normalized by cell number. One-way ANOVA ($F_{2,9}$ = 10.22, $p \leq 0.0048$), Dunnett's multiple comparisons test; $*p \leq 0.05$, $**p \leq 0.003$.

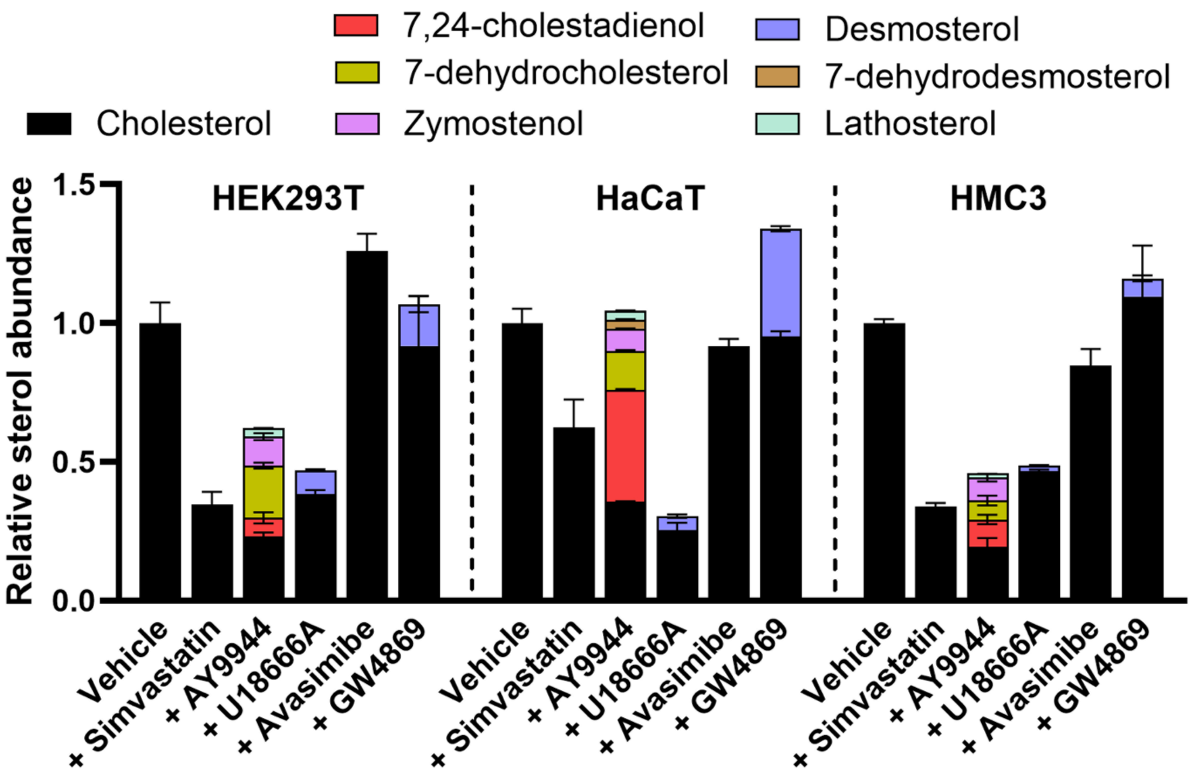


**Figure S3. Pharmacological inhibition of cholesterol metabolism shifts sterol profiles in three distinct cell types. Related to Figure 1.**

GC-MS analyses of HEK293T, HaCaT, and HMC3 after administration of small molecule inhibitors of cholesterol biosynthesis demonstrates a reduction in cholesterol and accumulation of sterol intermediates (mean ± SEM; n = 2 biological replicates from 2 independent experiments).

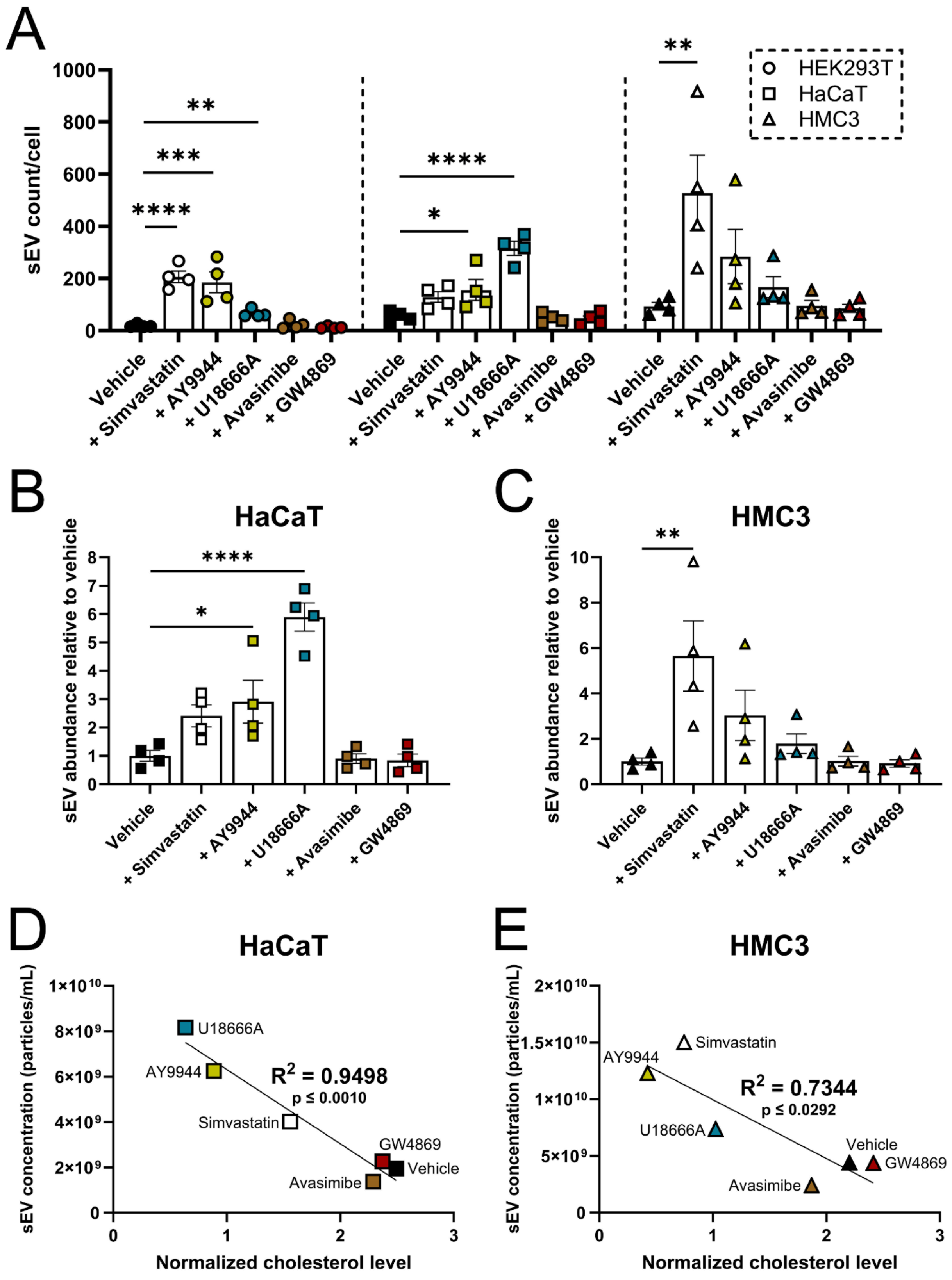

A
B
C
D
E
sEV count/cell
HEK293T
HaCaT
HMC3
Vehicle
+ Simvastatin
+ AY9944
+ U18666A
+ Avasimibe
+ GW4869
HaCaT
HMC3
sEV abundance relative to vehicle
sEV concentration (particles/mL)
Normalized cholesterol level
$R^2$ = 0.9498
p ≤ 0.0010
$R^2$ = 0.7344
p ≤ 0.0292

**Figure S4. Impaired cholesterol biosynthesis increases sEV release. Related to Figure 1.**

(A) sEV release is increased in HEK293T, HaCaT, and HMC3 cells following incubation with small molecule inhibitors of cholesterol biosynthesis (mean ± SEM; n = 4 biological replicates from 4 independent experiments). sEVs are normalized to cell number. One-way ANOVA (HEK293T: $F_{5,18} = 20.51$, $p < 0.0001$; HaCaT: $F_{5,18} = 21.25$, $p < 0.0001$; HMC3: $F_{5,18} = 5.308$, $p \leq 0.0036$), Dunnett's multiple comparisons test; *p ≤ 0.05, **p ≤ 0.01, ***p ≤ 0.001, ****p < 0.0001.

(B) Quantification of HaCaT sEV release in cholesterol metabolism inhibited cells relative to vehicle (mean ± SEM; n = 4 biological replicates from 4 independent experiments). One-way ANOVA ($F_{5,18} = 21.25$, $p < 0.0001$), Dunnett's multiple comparisons test; *p ≤ 0.05, ****p < 0.0001.

(C) HMC3 sEV release in cholesterol-targeted cells relative to vehicle treatment (mean ± SEM; n = 4 biological replicates from 4 independent experiments). One-way ANOVA ($F_{5,18} = 5.308$, $p \leq 0.0036$), Dunnett's multiple comparisons test; **p ≤ 0.005.

(D) Regression analysis shows a significant negative relationship between sEV secretion and cellular cholesterol levels in HaCaT (n = 6). Linear regression ($F_{1,4} = 75.63$, $R^2 = 0.9498$, $p \leq 0.001$)

(E) In HMC3 cells, regression analysis reveals a significant negative correlation between sEV release and cellular cholesterol levels (n = 6). Linear regression ($F_{1,4} = 11.06$, $R^2 = 0.7344$, $p \leq 0.0292$).

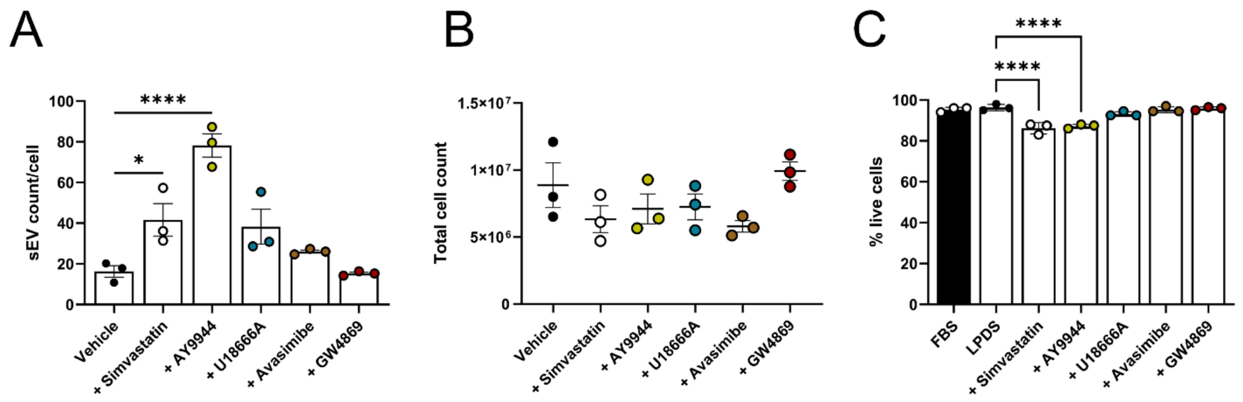


**Figure S5. Increased release of sEVs in response to cholesterol depletion is not time dependent. Related to Figure 1.**

(A) Increased sEV release in HEK293T cells is observed 24 h after treatment with simvastatin or AY9944 (mean ± SEM, n = 3 biological replicates from 3 independent experiments). sEVs are normalized to cell number. One way ANOVA ($F_{5,12}$ = 18.31, $p < 0.0001$), Dunnett's multiple comparisons test; *$p \leq 0.05$, ****$p < 0.0001$.

(B) Pharmacological inhibition of cholesterol biosynthesis for 24 h did not impact cell numbers (mean ± SEM; n = 3 biological replicates from 3 independent experiments).

(C) HEK293T cell viability was mildly impacted by cholesterol synthesis inhibition (mean ± SEM; n = 3 biological replicates from 3 independent experiments). One-way ANOVA ($F_{6,14}$ = 23.65, $p < 0.0001$), Dunnett's multiple comparisons test; ****$p <$ 0.0001.

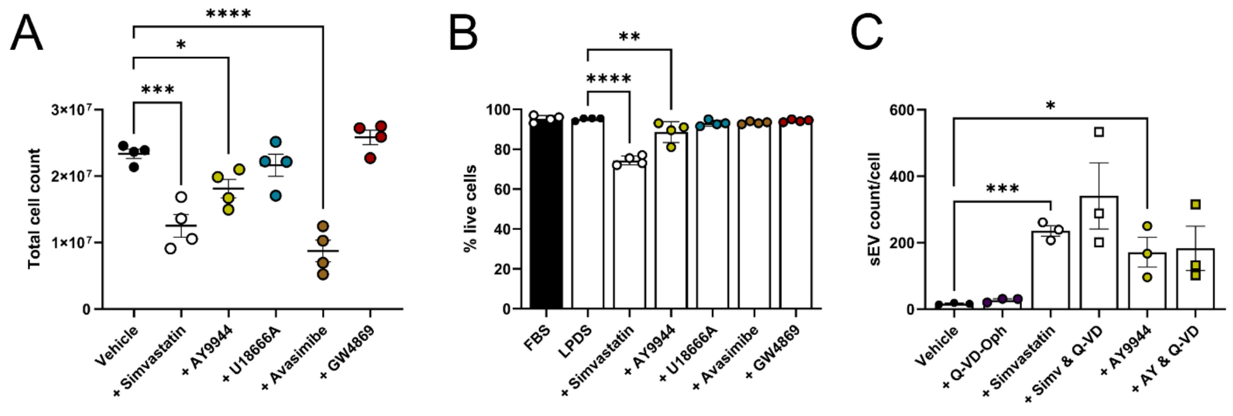


**Figure S6. Cholesterol biosynthesis inhibition does not cause increased release of apoptotic bodies. Related to Figure 1.**

(A) After 48 h cholesterol synthesis inhibition in HEK293T cells, simvastatin, AY9944, and avasimibe treatment reduced the total number of cells (mean ± SEM; n = 4 biological replicates from 4 independent experiments). One-way ANOVA ($F_{5,18}$ = 21.73, $p < 0.0001$). Dunnett's multiple comparisons test; *p ≤ 0.05, ***p ≤ 0.001, ****p < 0.0001.

(B) 48 h simvastatin and AY9944 treatment had a mild effect on cell viability (mean ± SEM; n = 4 biological replicates from 4 independent experiments). One-way ANOVA ($F_{6,21}$ = 39.51, $p < 0.0001$), Dunnett's multiple comparisons test; **p ≤ 0.005, ****p < 0.0001.

(C) Inhibition of apoptosis with the pan-caspase inhibitor Q-VD-Oph did not reduce sEV release following cholesterol synthesis inhibition by simvastatin or AY9944 (mean ± SEM; n = 3 biological replicates from 3 independent experiments). sEVs are normalized to cell number. One-way ANOVA ($F_{5,12}$ = 5.596, $p \leq 0.0069$). Unpaired t-tests (Vehicle vs Simvastatin: $t_4$ = 13.91, ***p ≤ 0.0002; Vehicle vs AY9944: $t_4$ = 3.477, *p ≤ 0.05).

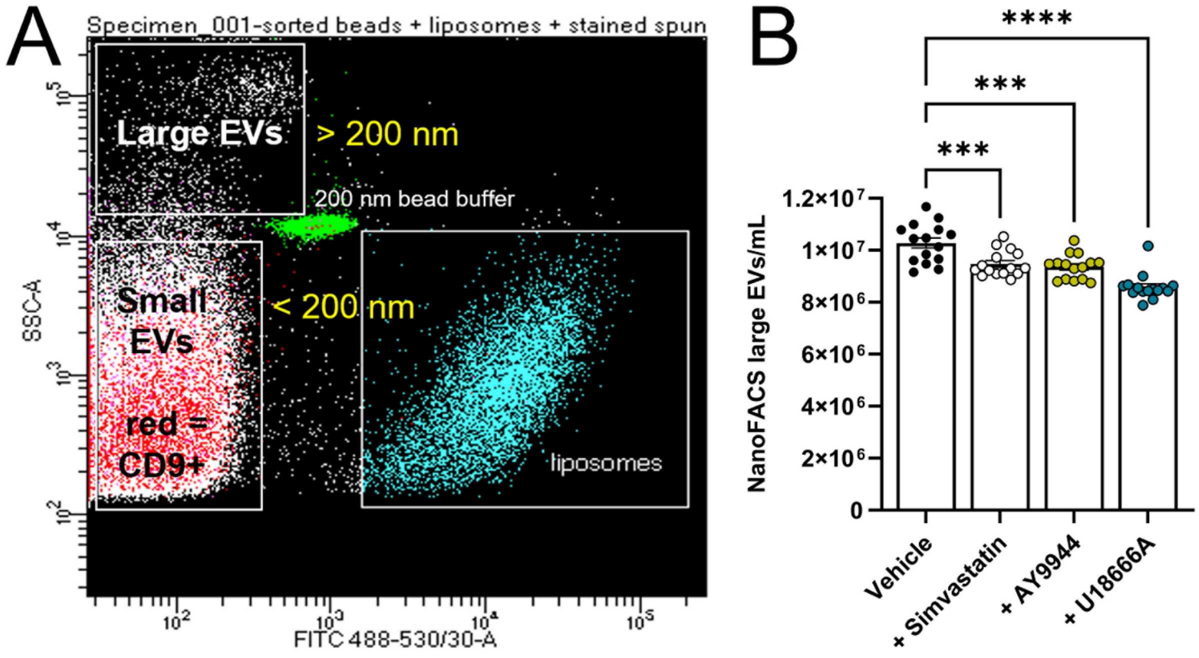


**Figure S7. Large EVs are reduced in media upon cholesterol depletion. Related to Figure 2.**

(A) Representative NanoFACS gating strategy for vesicle size showing CD9-positive vesicles from plasma for large EVs (200 nm – 900 nm), small EVs (<200nm) from plasma, and ~200 nm-sized fluorescently tagged liposomes.

(B) NanoFACS analyses of large (200 nm – 900 nm) EVs measured directly from media of treated HEK293T cells (mean ± SEM; n = 5 technical replicates/3 biological replicates from 3 independent experiments). One-way ANOVA ($F_{3,56} = 22.09$, $p < 0.0001$), Dunnett's multiple comparisons test; ***$p \leq 0.001$; ****$p < 0.0001$.

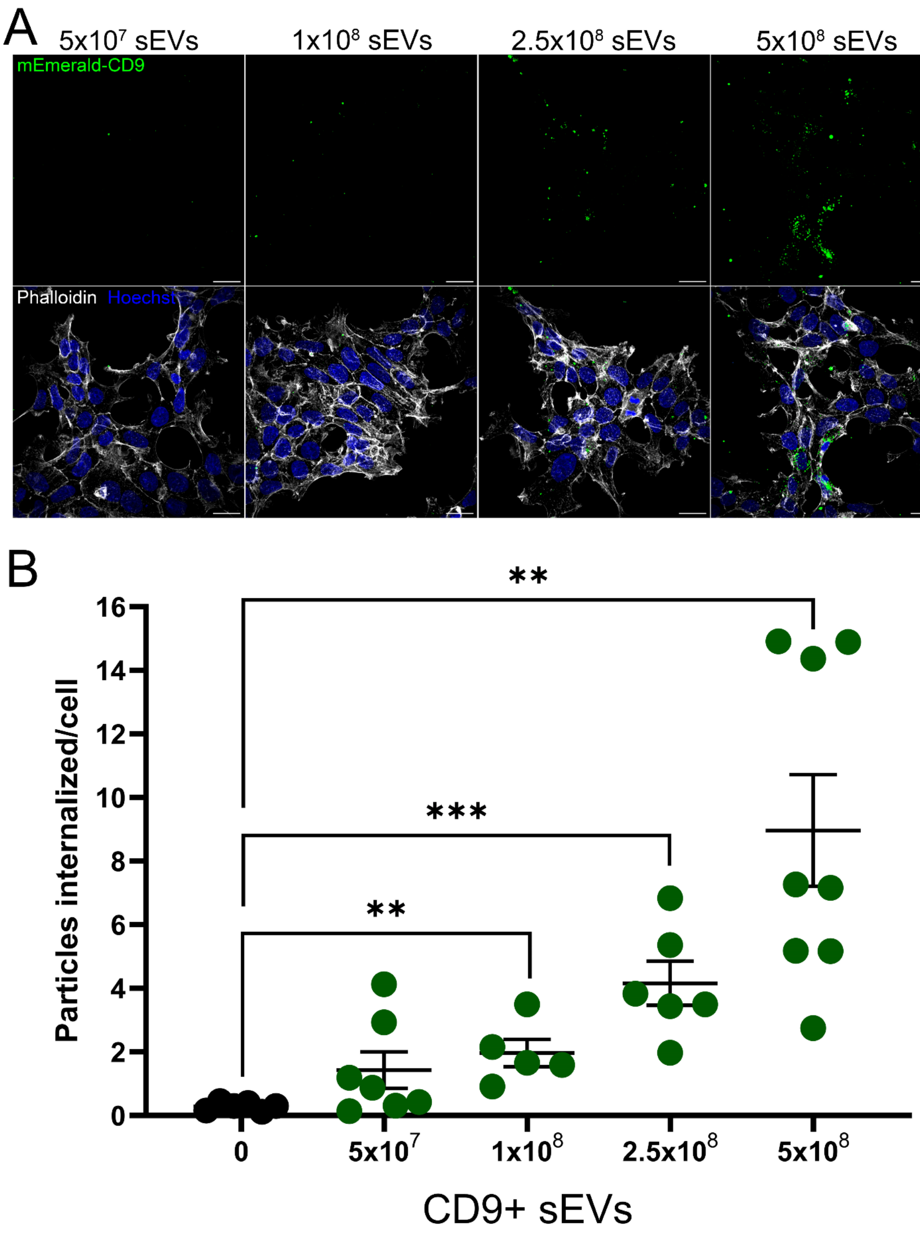


**Figure S8. Internalization of CD9-positive sEVs by recipient cells is dose-dependent. Related to Figure 4.**

(A) Representative images of recipient cell internalization of fluorescently tagged sEVs from HEK293T$^{mEmerald\text{-}CD9}$ cells occurs in a dose-dependent manner. The actin cytoskeleton is visualized with phalloidin. Scale bar, 25 µm.

(B) Quantification of mEmerald-CD9 sEV internalization in HEK293T cells (mean ± SEM; n = 5-8 images from 2 independent experiments). One-way ANOVA ($F_{2,27}$ = 11.31, $p < 0.0001$). Unpaired t-tests (0 vs 1x10$^8$ sEVs: $t_9$ = 4.248, **$p \leq 0.005$; 0 vs 2.5x10$^8$ sEVs: $t_{10}$ = 5.554, ***$p \leq 0.0002$; 0 vs 5x10$^8$ sEVs: $t_{12}$ = 4.228, **$p \leq 0.005$).

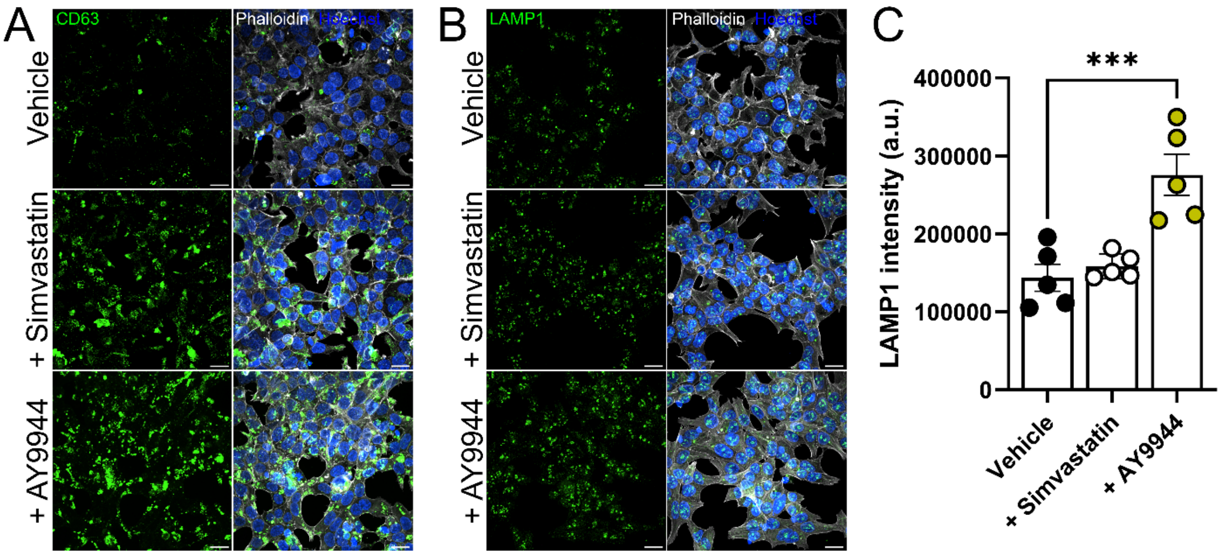


**Figure S9. Cholesterol deficiency results in increased expression of endosomal sorting markers. Related to Figure 5.**

(A) Representative images of CD63 in HEK293T cells show increased expression levels after simvastatin or AY9944 treatments. Scale bar, 25 μm.

(B) Representative images of LAMP1 in HEK293T cells demonstrate perinuclear lysosomal localization. Scale bar, 25 μm.

(C) Quantification of LAMP1 expression demonstrates an increase after AY9944 treatment compared to vehicle (mean ± SEM; n = 5). One-way ANOVA ($F_{2,12}$ = 14.97, $p \leq 0.0005$), Dunnett's multiple comparisons test; ***$p \leq 0.001$.

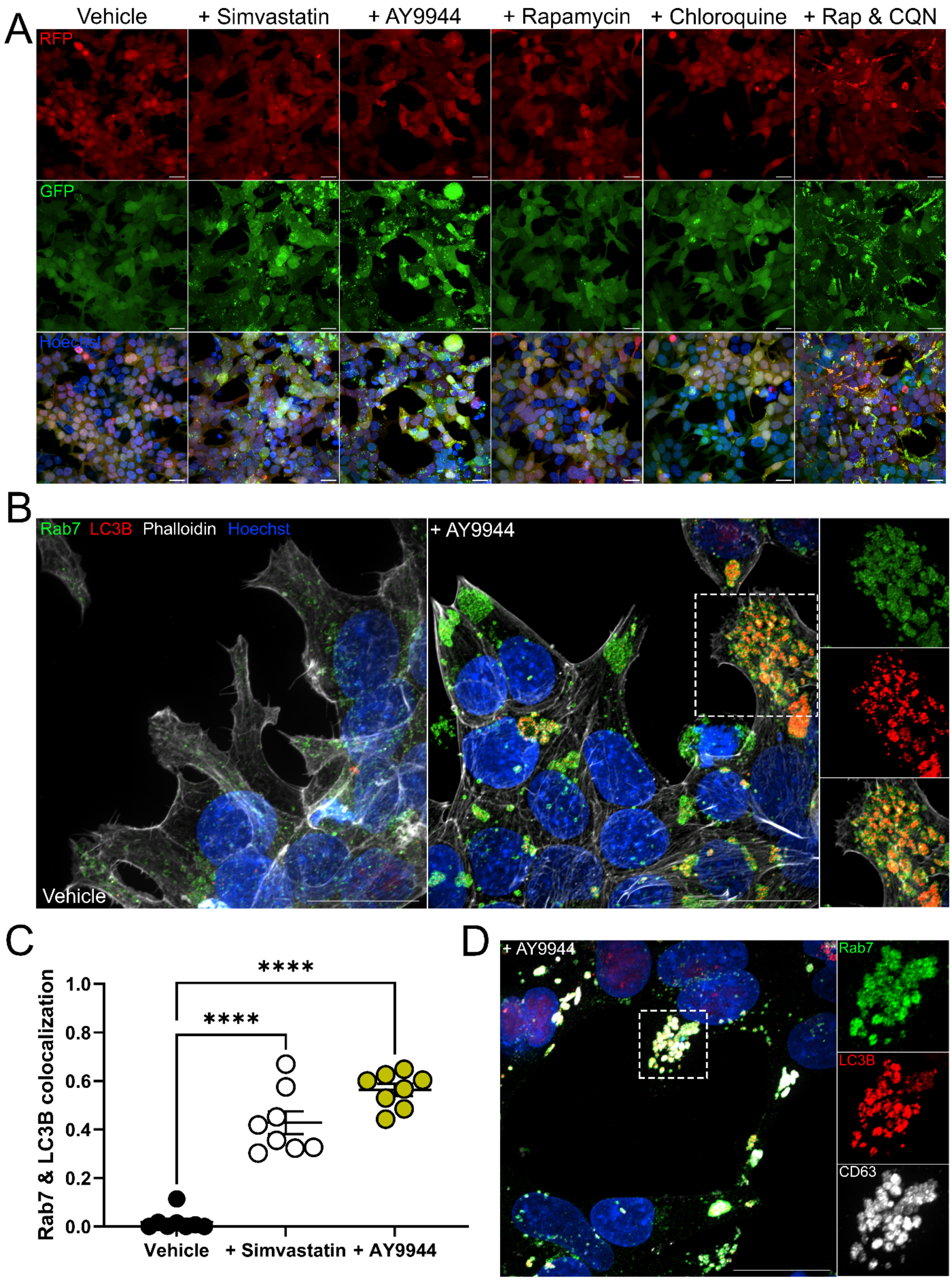

A
Vehicle
+ Simvastatin
+ AY9944
+ Rapamycin
+ Chloroquine
+ Rap & CQN
RFP
GFP
Hoechst
B
Rab7
LC3B
Phalloidin
Hoechst
Vehicle
+ AY9944
C
Rab7 & LC3B colocalization
1.0
0.8
0.6
0.4
0.2
0.0
****
****
Vehicle
+ Simvastatin
+ AY9944
D
+ AY9944
Rab7
LC3B
CD63

**Figure S10. Autophagosomes are directed toward late endosomes during cholesterol depletion. Related to Figure 6.**

(A) Representative images from HEK293T$^{GFP\text{-}LC3\text{-}RFP\text{-}LC3\Delta G}$ autophagic flux reporter cells demonstrate reduced autophagic flux (more GFP/RFP) after simvastatin, AY9944, chloroquine, or rapamycin & chloroquine (Rap & CQN) treatments. Scale bar, 25 μm.

(B) Representative images of HEK293T$^{LC3B\text{-}mCherry}$ cells reveal Rab7 and LC3B colocalization following AY9944 treatment. Scale bar, 25 μm.

(C) Colocalization of Rab7 and LC3B is increased by simvastatin or AY9944 treatment in HEK293T$^{LC3B\text{-}mCherry}$ cells (mean ± SEM; n = 8 images collected from 2 independent experiments). One-way ANOVA ($F_{2,21} = 79.46$, $p < 0.0001$), Dunnett's multiple comparisons test; ****p < 0.0001.

(D) Representative image of Rab7 and CD63 expression in HEK293T$^{LC3B\text{-}mCherry}$ cells demonstrates colocalization of Rab7, CD63, and LC3B after AY9944 treatment. Scale bar, 25 μm.

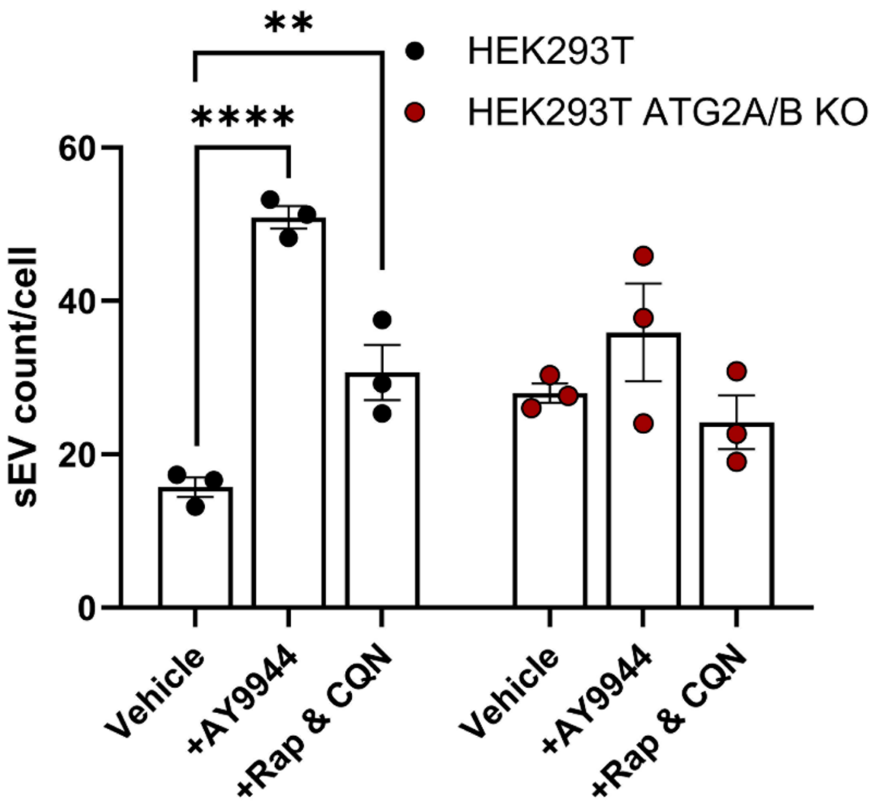


**Figure S11. Increased sEV secretion in response to cholesterol biosynthesis inhibition requires autophagic signaling. Related to Figure 7.**

Analyses of sEVs (normalized to cell counts) collected from HEK293T and $HEK293T^{ATG2A/B\ KO}$ cells (mean ± SEM; n = 3 biological replicates from 3 independent experiments). Two-way ANOVA (Treatment effect: $F_{2,12} = 21.1$, $p < 0.0001$; Interaction effect: $F_{2,12} = 8.195$, $p \leq 0.0057$). Sidak's multiple comparisons test (**$p \leq 0.01$, ****$p < 0.0001$).
