## Supplemental Table S1 for "Cholesterol deficiency directs autophagy-dependent secretion of extracellular vesicles"

| sEV Surface Marker | | Vehicle | | Simvastatin | | | AY9944 | | | U18666A | | | Avasimibe | | | GW4869 | | |
|---|---|---|---|---|---|---|---|---|---|---|---|---|---|---|---|---|---|---|
| **Full Name** | **Abrev Name** | **Intensity** | **±SEM** | **Intensity** | **±SEM** | **Exp** | **Intensity** | **±SEM** | **Exp** | **Intensity** | **±SEM** | **Exp** | **Intensity** | **±SEM** | **Exp** | **Intensity** | **±SEM** | **Exp** |
| Cluster of differentiation 3 | CD3 | 591.5 | 103.1 | 882.3 | 208.5 | 1.5 | **2447.8 *** | **876.5** | **4.1** | 914.6 | 120.3 | 1.5 | **125.4 *** | **18.8** | **0.2** | 445.0 | 41.7 | 0.8 |
| Major histocampatibility complex class II (MHC II) | HLA-DR,DP,DQ | 419.6 | 166.2 | 185.2 | 50.1 | 0.4 | 154.2 | 12.3 | 0.4 | 717.0 | 177.0 | 1.7 | 114.1 | 11.0 | 0.3 | 204.7 | 56.7 | 0.5 |
| Neural cell adhesion molecule 1 | CD56 | 1434.1 | 168.1 | **14663.2 **** | **5181.1** | **10.2** | **6960.4 **** | **1710.6** | **4.9** | **20280.7 ***** | **3557.9** | **14.1** | 1030.9 | 24.3 | 0.7 | 1108.3 | 59.3 | 0.8 |
| Interleukin-2 receptor alpha chain | CD25 | 377.2 | 142.8 | **66.2 *** | **13.2** | **0.2** | **55.9 *** | **6.8** | **0.1** | 98.2 | 5.0 | 0.3 | 117.8 | 43.7 | 0.3 | 380.0 | 44.7 | 1.0 |
| Integrin alpha-5 | CD49e | 2124.9 | 1155.4 | 3922.8 | 3297.1 | 1.8 | 1855.3 | 1439.8 | 0.9 | 5824.8 | 4851.7 | 2.7 | 5665.9 | 470.8 | 2.7 | 328.3 | 21.2 | 0.2 |
| Receptor tyrosine kinase-like orphan receptor 1 | ROR1 | 943.6 | 171.2 | **5671.4 ******* | **977.9** | **6.0** | **2587.6 **** | **329.3** | **2.7** | **2910.4 ***** | **259.8** | **3.1** | 933.3 | 122.9 | 1.0 | 1248.0 | 102.3 | 1.3 |
| Cluster of differentiation 9 (Tetraspanin 29) | CD9 | 33435.1 | 2384.3 | 39514.0 | 9062.7 | 1.2 | 27106.7 | 1707.6 | 0.8 | 44017.0 | 4920.1 | 1.3 | 33721.2 | 2841.0 | 1.0 | 34086.7 | 594.5 | 1.0 |
| Stage-specific embryonic antigen-4 | SSEA-4 | 1671.4 | 370.7 | **487.3 *** | **166.0** | **0.3** | 936.5 | 259.5 | 0.6 | 868.8 | 209.2 | 0.5 | 2069.9 | 217.7 | 1.2 | 1155.7 | 275.0 | 0.7 |
| Major histocampatibility complex class I (MHC I) | HLA-A,B,C | 369.4 | 94.2 | 459.9 | 125.5 | 1.2 | 441.9 | 59.7 | 1.2 | 577.9 | 98.8 | 1.6 | 283.5 | 26.5 | 0.8 | 343.3 | 128.8 | 0.9 |
| Cluster of differentiation 63 (Tetraspanin 30) | CD63 | 33723.8 | 2251.7 | 37802.5 | 7865.4 | 1.1 | **24685.8 *** | **1469.0** | **0.7** | 41853.0 | 5081.7 | 1.2 | 29953.5 | 2435.9 | 0.9 | 34767.7 | 1120.1 | 1.0 |
| Cluster of differentiation 81 (Tetraspanin 28) | CD81 | 30913.3 | 1146.0 | 36209.1 | 8369.8 | 1.2 | **24428.1 **** | **1050.3** | **0.8** | 31538.3 | 2157.4 | 1.0 | 31550.9 | 1404.7 | 1.0 | 31435.3 | 669.9 | 1.0 |
| Melanoma-associdated chondroitin sulfate proteoglycan | MCSP | 413.3 | 132.2 | 198.1 | 24.5 | 0.5 | **118.9 *** | **10.4** | **0.3** | 433.6 | 30.7 | 1.0 | **88.6 *** | **12.1** | **0.2** | 329.3 | 28.9 | 0.8 |
| Melanoma cell adhesion molecule | CD146 | 2799.9 | 578.9 | 10656.0 | 4337.9 | 3.8 | **5421.3 *** | **853.8** | **1.9** | **18221.3 **** | **3855.9** | **6.5** | 981.8 | 157.5 | 0.4 | 2332.0 | 171.5 | 0.8 |
| Integrin alpha-IIb | CD41b | 1621.7 | 534.4 | 550.2 | 194.9 | 0.3 | **274.0 *** | **129.4** | **0.2** | 685.2 | 270.4 | 0.4 | 1124.3 | 143.3 | 0.7 | 599.7 | 255.7 | 0.4 |
| Cluster of differentiation 24 (Heat stable antigen) | CD24 | 2951.1 | 759.8 | 2850.2 | 497.8 | 1.0 | 1335.5 | 196.3 | 0.5 | **5024.5 *** | **314.2** | **1.7** | 1496.4 | 97.5 | 0.5 | 1192.0 | 330.0 | 0.4 |
| Epithelial cell adhesion molecule (EpCAM) | CD326 | 5130.7 | 1242.8 | 9348.4 | 3690.5 | 1.8 | 4907.6 | 465.5 | 1.0 | **13297.6 *** | **3139.7** | **2.6** | 3522.5 | 333.2 | 0.7 | 7225.3 | 141.6 | 1.4 |
| Cluster of differentiation 133/1 (Prominin-1) | CD133/1 | 4655.0 | 1598.7 | 5859.3 | 884.6 | 1.3 | 3650.5 | 379.7 | 0.8 | 7648.0 | 1016.6 | 1.6 | 919.1 | 159.0 | 0.2 | **10057.3 *** | **649.7** | **2.2** |
| Integrin β1 | CD29 | 22692.9 | 3516.7 | 52532.2 | 15253.8 | 2.3 | 26584.5 | 8069.2 | 1.2 | **60930.0 ***** | **5925.7** | **2.7** | **10035.9 *** | **1111.1** | **0.4** | 21674.3 | 2837.5 | 1.0 |
| Protein tyrosine phosphatase receptor type C (PTPRC) | CD45 | 409.7 | 83.7 | 391.7 | 61.3 | 1.0 | 654.9 | 141.1 | 1.6 | 545.9 | 41.0 | 1.3 | 138.3 | 20.8 | 0.3 | 430.7 | 34.1 | 1.1 |

Intensity = mean of median intensity values from independent experiments; ±SEM = standard error of mean; Exp = expression level relative to vehicle (treatment avg intensity/vehicle avg intensity); Unpaired t-test compared to vehicle: *$p \leq 0.05$, **$p \leq 0.01$, ***$p \leq 0.001$, ****$p < 0.0001$
